# Reverse Engineering of Feedforward Cortical-Hippocampal Microcircuits for Modelling Neural Network Function and Dysfunction

**DOI:** 10.1101/2023.06.26.546556

**Authors:** Katrine Sjaastad Hanssen, Nicolai Winter-Hjelm, Salome Nora Niethammer, Asgeir Kobro-Flatmoen, Menno P. Witter, Axel Sandvig, Ioanna Sandvig

## Abstract

Engineered biological neural networks are indispensable models for investigation of neural function and dysfunction from the subcellular to the network level. Notably, advanced neuro-engineering approaches are of significant interest for their potential to replicate the topological and functional organization of brain networks. In this study, we reverse engineered feed-forward neural networks of primary cortical and hippocampal neurons, using a custom-designed multinodal microfluidic device with Tesla valve inspired microtunnels. By interfacing this device with nanoporous microelectrodes, we show that the reverse engineered multinodal neural networks exhibit capacity for both segregated and integrated functional activity, mimicking brain network dynamics. To advocate the broader applicability of our model system, we induced localized perturbations with amyloid beta to study the impact of pathology on network functionality. Additionally, we demonstrate long-term culturing of subregion- and layer specific neurons extracted from the entorhinal cortex and hippocampus of adult Alzheimer’
ss-model mice and rats. Our results thus highlight the potential of our approach for reverse engineering of anatomically relevant multinodal neural networks to study dynamic structure-function relationships in both healthy and pathological conditions.

## Introduction

Engineered biological neural networks facilitate the study of dynamic properties of network structure and function at the microscale (i.e., subcellular and cellular level) and mesoscale (i.e., network level), in highly controllable experimental conditions. There is compelling evidence that neurons *in vitro* maintain their inherent self-organizing properties and over time form neural networks with complex structure and function, thus recapitulating fundamental behaviour of brain networks (1–4). As such, engineered neural networks offer a complementary approach to *in vivo* studies for investigations of neural structure and function from the subcellular to the network level. In recent years, several methodological and technological developments have enabled advanced neural network modelling *in vitro* using microfluidic devices to engineer multinodal neural networks (5, 6). However, most studies to date are typically limited to using two-nodal mi-crofluidic devices (7, 8). Thus, the potential of such neuro-engineering approaches to recapitulate anatomically relevant network configurations is still underutilized. Additionally, the relevant models tend to be limited in terms of their reproducibility, scalability, and adaptability to support multinodal, directional topological configurations resembling *in vivo* neural assemblies.

A key determinant for establishing anatomically relevant neural network configurations *in vitro* is facilitation and control of multinodality. Neural assemblies in the brain are structured into highly specialized functional regions connected through precise unidirectional axonal pathways, aiding feedforward and/or feedback signal propagation between the nodes (9). During neural development in the brain, directional projection sequences are established by finely tuned spatiotemporally regulated molecular and chemical cues determining axon growth and guidance (10–12). Since such gradients are absent in an *in vitro* microenvironment, alternative approaches are required to facilitate establishment of directional projections between the neural nodes. Several recent studies, including studies from our group, have demonstrated that directional axonal outgrowth between interconnected nodes can be achieved and controlled by embedding geometrical constraints within the internodal microtunnel architecture in microfluidic interfaces (13–20). This approach supports control of connectivity between multiple neural subpopulations with high precision, rendering them particularly suitable for recapitulating anatomically relevant neural networks (21–24). By interfacing this type of microfluidic devices with microelectrode arrays (MEAs), emerging structure-function dynamics can also be studied electrophysiologically (25). Previous studies suggest that containment of neural networks into modular configurations can promote the development of a broader repertoire of activity dynamics compared to unstructured networks, akin to complex information processing seen *in vivo* (22, 26). For instance, we and others have shown that structurally coupled neural networks exhibit functional asymmetry, with independently regulated information processing within the interconnected nodes (i.e., intranodal), as well as controlled information flow between them (i.e., internodal) (20, 21, 27, 28).

A relevant neural network configuration to study by using such interfaces is the highly interconnected entorhinal-hippocampal network, a brain area important for processing and storage of memories (29). The entorhinal-hippocampal network consists of multiple subregion specific cell types connected through axonal projections. Multinodal microfluidic devices offer an opportunity to reconstruct this network, and to study ongoing structural and functional dynamics in a highly controlled microenvironment. Furthermore, the entorhinal-hippocampal network is particularly vulnerable to Alzheimer’s disease (AD), evidenced by early accumulation of tau pathology (30), amyloid beta plaque load (31), altered neuronal activity such as seizure-like hyperactivity (32), and major neuronal (33, 34) and synaptic loss (35, 36). Thus, this network is of high interest to study at preclinical stages of AD. We and others have demonstrated that adult entorhinal and hippocampal subregion specific cells from AD model mice are able to reconnect in culture (37–39). We also recently reported a method for long-term (>2 months) culturing and recording of lateral entorhinal cortex layer II (LEC LII) neurons from AD model animals *in vitro* (39).

In the present study, we demonstrate reverse engineering of anatomically relevant biological neural networks, including networks of layer specific neurons derived from adult AD model animals, using custom-designed microfluidic MEA interfaces. By incorporating Tesla valve inspired microtunnels between nodes, we could structure multinodal neural networks with controllable feedforward connectivity, and record spontaneously evoked and stimulation induced signal propagation between the nodes. This engineering approach supports the longitudinal study of dynamic structure-function relationships in both healthy and diseased neural networks with high temporal and spatial precision. We furthermore demonstrate the applicability of this model system for selectively inducing localized perturbations with human amyloid beta (A*β*) fragments, allowing us to study the spread and impact of AD relevant pathology on network functionality. Moreover, we demonstrate long-term culturing of region and layer specific neurons extracted from adult AD-model mice and rats on these platforms in an anatomically relevant configuration. We show that these adult neurons re-form structural connections and develop spontaneous electrophysiological spiking activity after 15 days *in vitro* (DIV). Taken together, our results demonstrate that our reverse engineering approach is highly relevant for advanced preclinical modelling of neural network function and dysfunction, including neurodegeneration.

## Results

### Feedforward Cortical-Hippocampal Networks Are Readily Established in The Four-Nodal Microdevices

In this study, we reverse engineered multinodal rodent cortical-hippocampal neural networks with controllable unidirectional connectivity using custom-designed mi-crofluidic devices interfaced with microelectrode arrays. Neurons within the six four-nodal devices self-organized into highly complex networks by 14 DIV (**Figure 1A**). At this stage, electrophysiological recordings furthermore indicated that spontaneous neural activity had started transitioning from immature tonic firing to synchronous bursting within the nodes, consistent with previous findings (**Figure 1B**) (1, 3, 20, 40, 41).

**Figure 1.**
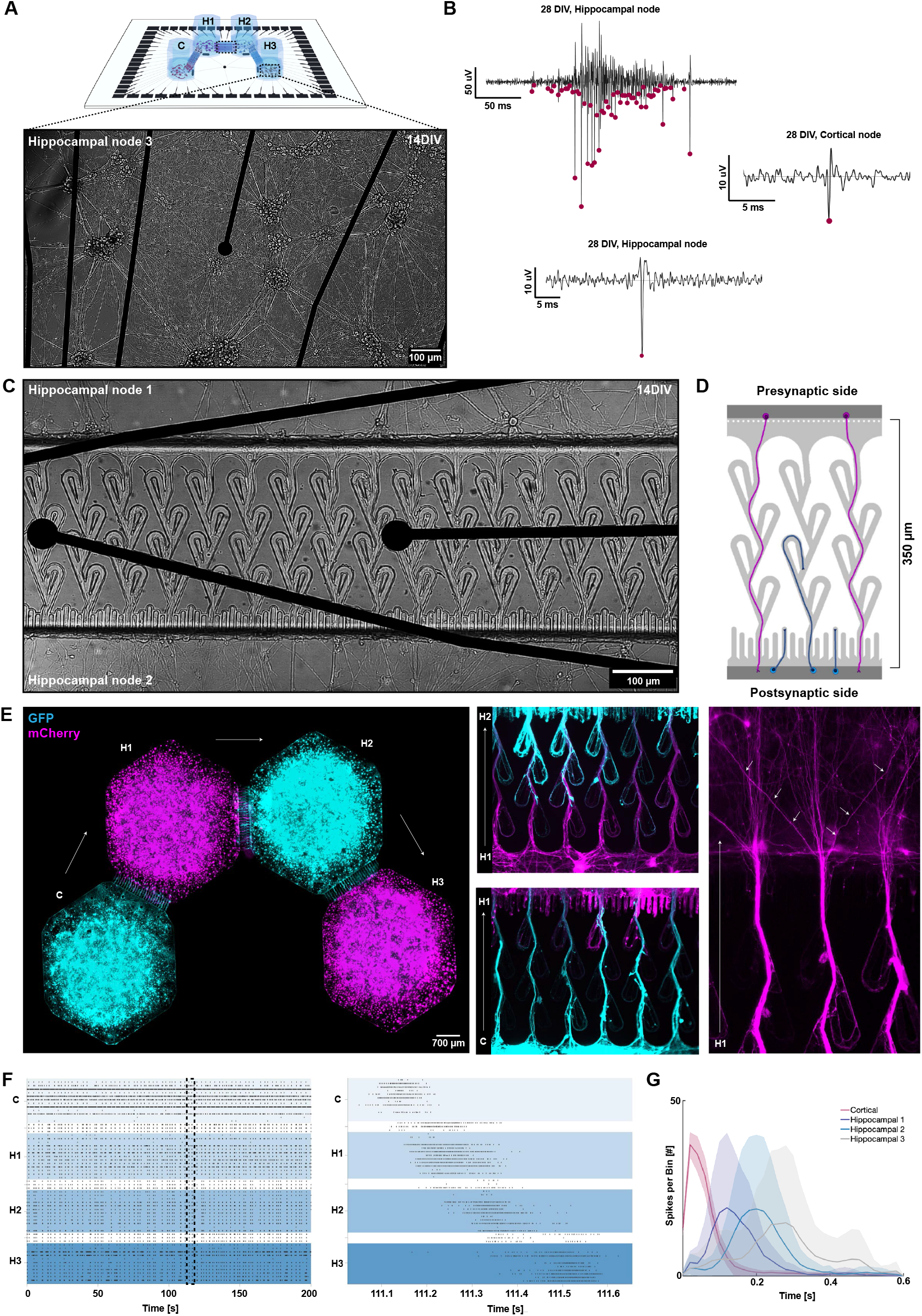
Feedforward connectivity establishment and activity propagation in the four-nodal cortical-hippocampal networks. **(A.)** Representative micrograph of a hippocampal node at 14 DIV. **(B.)** Voltage traces showing a burst detected within one of the hippocampal nodes at 28 DIV, as well as representative examples of spikes detected within the cortical and hippocampal nodes. Pink specks represent spikes detected by the PTSD algorithm. **(C.)** Micrograph showing neurites crossing the Tesla valve inspired microtunnels already after 14 DIV. **(D.)** Graphical illustration of how the Tesla valve design promotes neurite outgrowth from the purple cell population across the microtunnels, whereas neurites from the blue cell population get redirected to the chamber from which they originated by the Tesla loops or trapped in the saw-tooths. **(E.)** Micrograph of a four-nodal network with neurons expressing either GFP (node C and H2, cyan) or mCherry (nodes H1 and H3, magenta). The higher magnification images show how neurites from the presynaptic sides (i.e. from C to H1 or H1 to H2) fill up the microtunnels and defasciculate into the opposite nodes, while the neurites from the postsynaptic sides are rerouted back to their node of origin by the Tesla valves. **(F.)** Representative raster plot showing the complex network dynamics emerging in the four-nodal networks. A fraction of the network bursts were also clearly spreading from the cortical node throughout all four nodes, as represented in the zoomed-in graph. **(G.)** Peristimulus time histogram displaying the average response of each of the four nodes following repeated (n=60) stimulations of the cortical node for one network at 28 DIV. The stimulations clearly induced feedforward activity propagating throughout all four nodes. DIV: Days *In Vitro*, C: Cortical node, H1-3: Hippocampal node 1-3.

The microfluidic tunnels connecting the individual nodes were designed to promote unidirectional structural connectivity. Tesla valves were included to redirect axons from the postsynaptic side back to their node of origin, while sawtooths were included on the postsynaptic side to misguide axons growing towards the microtunnel inlets (**Figures 1C** and **1D**). To verify unidirectionality, neurons were transduced with AAV viruses for ubiquitous expression of either GFP (nodes 1 and 3) or mCherry (nodes 2 and 4). This verified that the design indeed guided neurites originating in the presynaptic node all the way through the microtunnels, while neurites from the postsynaptic node were rerouted back to their node of origin (**Figure 1E**).

Functionally, both spontaneously evoked and stimulation induced network bursts were used to assess the capacity of the networks for feedforward activity propagation between the four unidirectionally connected nodes. A fraction of the network bursts initiated in the cortical node, i.e., node 1, could be seen spreading in a feedforward manner throughout all four nodes already at 12 DIV (**Figure 1F**). Furthermore, electrical stimulations delivered to the cortical node initiated activity propagating sequentially along the four nodes (**Figure 1G**).

### Functional Integration Increases, While Functional Segregation is Maintained Over Time in Reverse Engineered Multinodal Microcircuits

The functional connectivity of the networks was evaluated using Pearson correlation and pairwise mutual information. Both intranodal and internodal correlation increased over time (**Figure 2A**). While some internodal correlation could be observed between nodes already at 12 DIV, integration of all neighboring nodes was not properly established until 20 DIV. At 28 DIV, a higher correlation was also seen between non-neighboring nodes, indicative of a higher network-wide synchronization at this point (20, 27, 28). Graph representations of the networks furthermore confirmed that the internodal correlation was clearly strongest between neighboring nodes, while the intranodal correlation on average was higher than the internodal correlation (**Figure 2B**).

**Figure 2.**
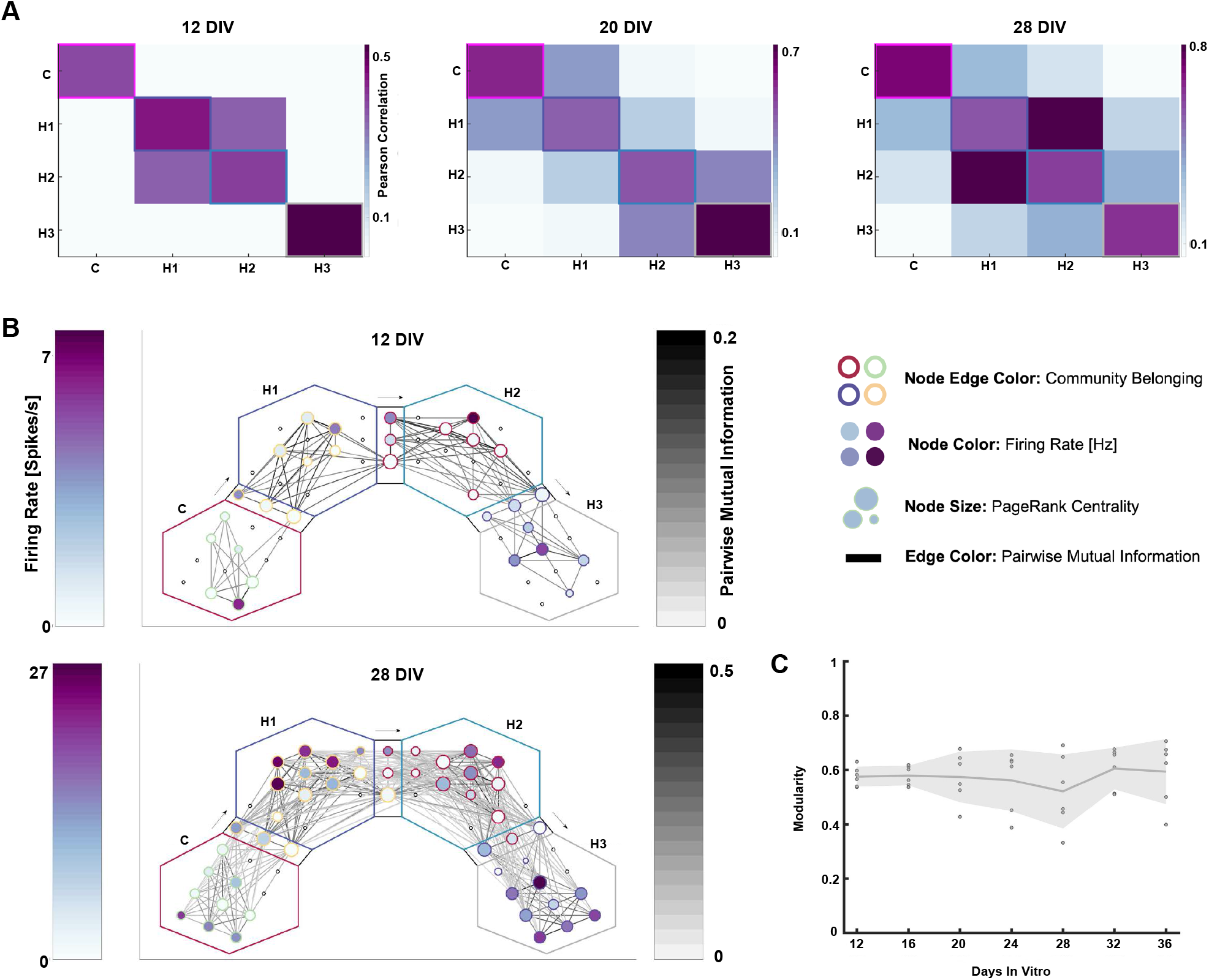
Functional connectivity of the cortical-hippocampal networks. **(A.)** Correlation matrices showing the temporal increase in functional connectivity of one representative network over time from 12 to 28 DIV. **(B.)** Representative graphs of a single network at 12 and 28 DIV displaying individual electrodes as nodes and their correlation as edges, i.e., lines connecting the nodes. Node color represents firing rate, node size pagerank centrality and the edge color around the nodes which community the node was classified as when the graph was analysed with the Louvain algorithm. As illustrated, the network can be clearly delineated into distinct communities, with higher intranodal than internodal functional connectivity. These communities are furthermore localized within the distinct nodes of the microfluidic interface. **(C.)** Modularity for the four-nodal networks, indicating a retained functional segregation over time.

Furthermore, the entire reverse-engineered cortical-hippocampal networks could be mathematically delineated into four distinct modules using the Louvain algorithm (**Figure 2B**). Each of the four detected communities were furthermore physically located within each of the four individual nodes of the microfluidic interface. Despite exhibiting higher internodal correlation with time, the median modularity value stayed close to 0.6 for all networks throughout the experimental period, demonstrating that the networks retained functional segregation of the four nodes (**Figure 2C**). The activity recorded by electrodes within the microtunnels also indicated a high pagerank centrality, high-lighting their importance for information transfer within the networks (**Figure 2B**). These results consequently confirmed that the networks matured into highly integrated multinodal networks with time, while still retaining high capacity for local information processing within the individual nodes.

### Cortical-Hippocampal Networks Exhibit Signs of Both Structural Maturation and Plasticity at 27 DIV

As we saw a high level of functional integration between the nodes beyond 20 DIV, we also evaluated the networks’ structural maturation using immunocytochemistry. To do so, we stained cells in each of the four nodes for both developmental and mature cytoskeletal and nucleic markers at 27 DIV. High levels of Growth-Associated Protein 43 (GAP43) were expressed in both the cortical and hippocampal nodes at 27 DIV (**Figure 3A**). This marker was used in combination with the cytoskeletal proteins Microtubule-Associated Protein 2 (MAP2) and *β*3-Tubulin, which are expressed from early axogenesis and onwards (42–44). As GAP43 is critical for neurodevelopment and growth cone migration, the high expression indicates that structural processes were still evolving beyond three weeks *in vitro* (45). Furthermore, both Neural Nuclear Protein (NeuN) and Neurofilament Heavy (NFH) have been found to be specific for mature neurons, and were used here to assess the maturity of the engineered networks (**Figure 3B**) (46, 47). These markers were used in combination with the glial marker GFAP (48). We found all these proteins to be prominently expressed at 27 DIV, indicating that the networks were reaching structural maturity. This is also consistent with the findings from the electrophysiology, showing highly integrated dynamics between the nodes at this time point (**Figure 2**).

**Figure 3.**
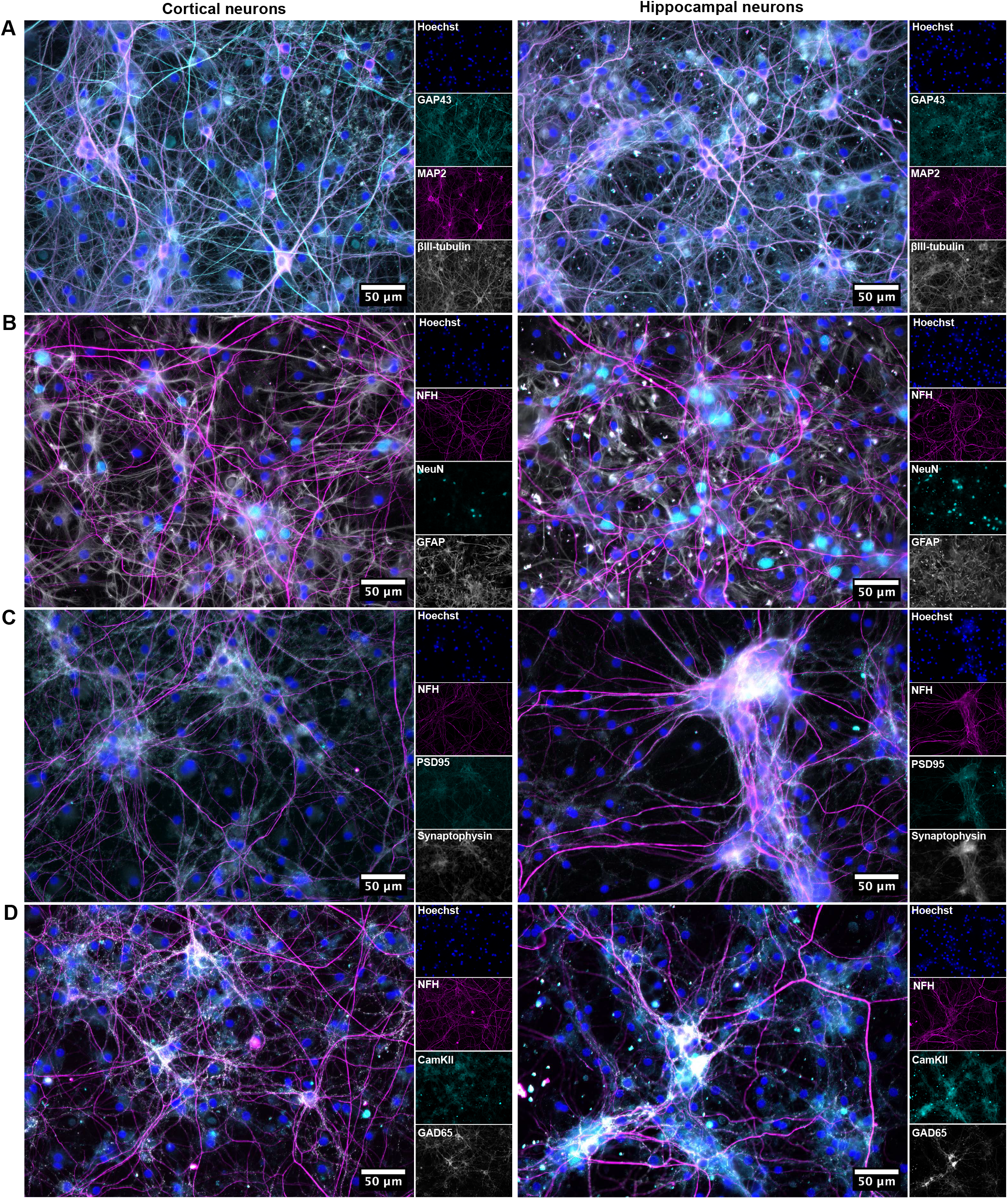
Immunocytochemistry of cortical-hippocampal networks. **(A.)** Both cortical and hippocampal neurons self-organized into complex networks, as indicated by the cytoskeletal marker proteins *β*3-Tubulin and MAP2. The networks still underwent some level of self-organization and maturation at 27 DIV, as indicated by GAP43, a protein critical for neurodevelopment and growth cone migration. **(B.)** The networks had reached a high level of structural maturity at 27 DIV, indicated by the mature markers Neurofilament Heavy (NFH) and NeuN. **(C.)** Colocalization of the pre- and postsynaptic markers synaptophysin and PSD95 indicated mature synapses. **(D.)** Expression of the glutamatergic and GABAergic markers CaMKII and GAD65 indicated the presence of both excitatory and inhibitory neurons in the networks.

Colocalization of the pre- and postsynaptic markers synapto-physin and PSD95 verified the presence of mature synaptic connections (**Figure 3C**) (49, 50). Furthermore, distinct expression of CaMKII and GAD65 verified the presence of both glutamatergic and GABAergic neurons, respectively (**Figure 3D**) (41, 51–53).

### Amyloid Beta Oligomerisation Remain Localized in Perturbed Nodes, but Weaken Overall Network Integration

To showcase the applicability of the model system for disease modelling, human A*β*-fragments were added to the cortical node of six four-nodal networks at 23 DIV. This perturbation led to a significant gradual increase in the number of A*β*-oligomers in the cortical node over time (**Figures 4A** and **4B**, p = 0.0286). Neither any of the hippocampal nodes, nor the unperturbed control networks, exhibited expression of such A*β*-oligomers during this period of time (**Figure 4C**).

**Figure 4.**
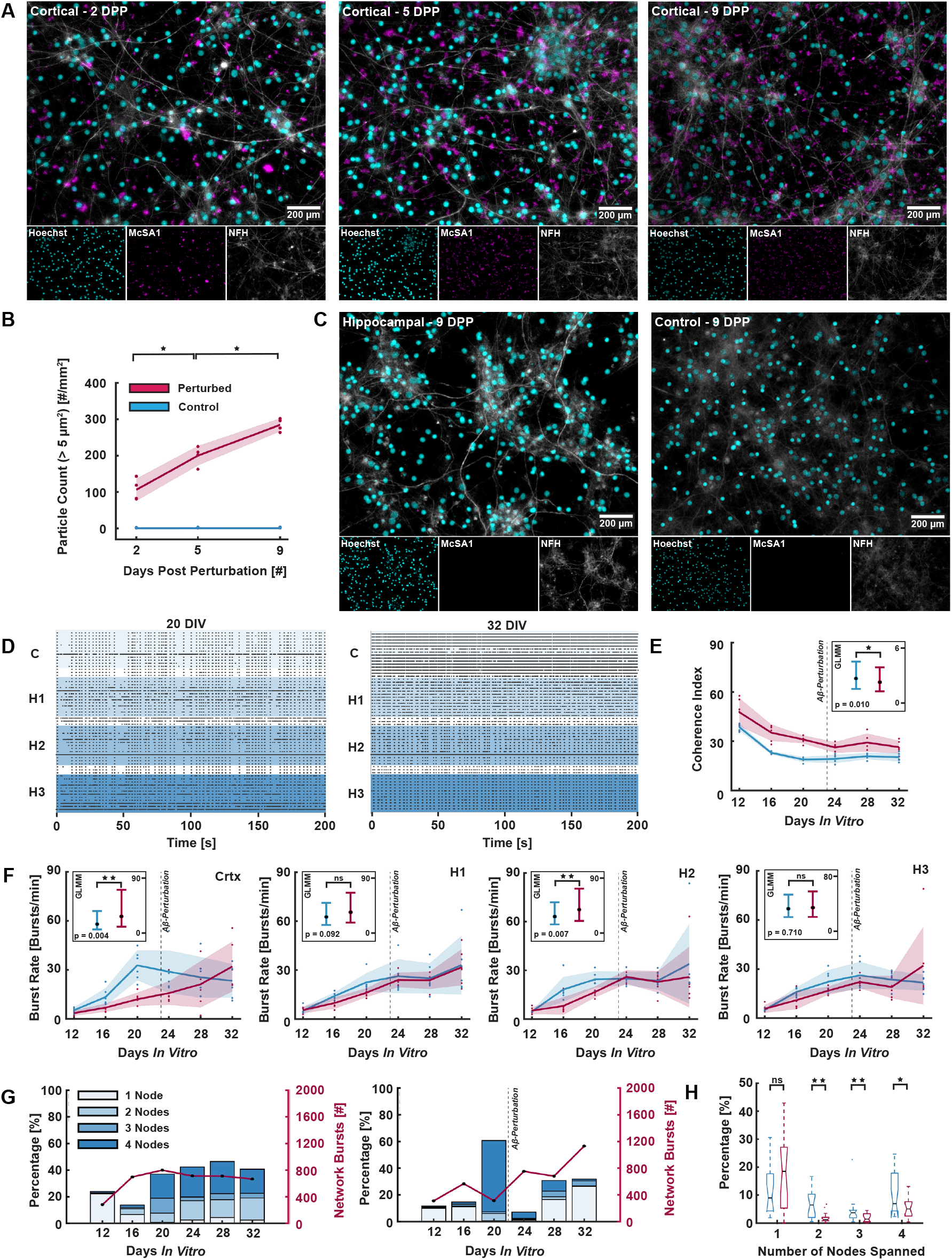
Impact of amyloid beta perturbations on network activity. **(A.)** The number of amyloid beta (A*β*) oligomers increased over time in the cortical node following perturbations, visualized using the antibody McSA1 staining for insoluble A*β*-aggregates. **(B.)** Such aggregates were however not seen in the hippocampal nodes, nor the unperturbed control networks at 2, 5 or 9 days post perturbation (DPP). **(C.)** The number of particles above 5 µm in diameter increased significantly between 2 - 5 DPP and 5 - 9 DPP (p = 0.0286 for both). **(D.)** The activity increased over time in particularly the cortical node following perturbations, as illustrated in the raster plots at 20 DIV (prior to perturbation) and 32 DIV (9 days after perturbation) for one representative network. **(E.)** The coherence index, indicating network synchrony, was found significantly lower for networks after perturbation (n = 6, 24 - 32 DIV) when compared to the perturbed networks prior to perturbation (n = 6, 12 - 20 DIV) and control networks (n = 6, 12 - 32 DIV) using GLMM estimated group averages with 95 % confidence intervals. **(F.)** The burst rate was found significantly higher in the perturbed networks of the cortical node and the second hippocampal node when evaluating the GLMM estimated group averages. The difference between the perturbed and unperturbed networks of the other two hippocampal nodes were however found to be non-significant. **(G.)** Histogram showing the percentage of spontaneously evoked network bursts initiated within the cortical population, and the number of nodes they spanned, for one representative control network and one perturbed network. The total number of network bursts initiated at each DIV is shown along the secondary y-axis. A significant change can be seen for the perturbed networks between DIV 20 (prior to perturbation) and DIV 24 (after perturbation). **(H.)** The percentage of network bursts initiated in the cortical node spanning 2, 3 and 4 nodes was on average significantly lower for the perturbed networks compared to the controls (p = 0.001, p = 0.005 and p = 0.035, respectively).

To examine whether this perturbation affected the electrophysiological behaviour of the networks, we further investigated the change in functional dynamics both within and across nodes of the six four-nodal networks interfaced with MEAs. Both perturbed and control networks transitioned to exhibiting gradually higher firing rates over time. The activity of the cortical node did however appear to transition towards a more hyperactive state after perturbation with A*β* (**Figure 4D**). Furthermore, the coherence index, measuring network synchrony, was found to be significantly lower in the perturbed networks when compared to the controls using Generalized Linear Mixed Effect Model (GLMM) estimated group averages (**Figure 4E**). In line with this, the burst rate was found to be significantly higher in the cortical node in the perturbed networks when assessing the GLMM estimated group averages (**Figure 4F**). While the burst rate was also found higher in the second hippocampal node, the difference between the perturbed and unperturbed networks was found non-significant in the first and third hippocampal nodes.

The percentage of network bursts initiated in the cortical node and spreading to the next one, two or three nodes increased significantly between 12 and 20 DIV for both controls and perturbed networks (**Figure 4G**). In the control networks, at 28 DIV, as much as 24.2 % of the network bursts propagated through all four nodes, compared to less than 2 % at 12 DIV. For comparison 4.5 %, 13.4 % and 4.4 % of the network bursts that were initiated in the cortical node propagated and stopped after the first, second and third node, respectively. This indicated establishment of more mature directional projection sequences between the nodes. The remaining percentages of network bursts were initiated within the second, third or fourth node. Similar trends were found for the other five control networks (**Figure S1**). In the perturbed networks, a sharp decline in the number of network bursts spanning beyond the cortical node was observed between 20 DIV (prior to perturbations) and 24 DIV (following perturbations). There were furthermore on average significantly fewer network bursts initiated in the cortical node spanning 2, 3 or 4 nodes in the perturbed networks in the recordings at 24, 28 and 32 DIV compared to the control networks (**Figure 4H**, p = 0.001, p = 0.005 and p = 0.035, respectively).

### Adult Entorhinal and Hippocampal AD Neurons Re– form Structural Networks and Exhibit Electrophysio-logical Activity

To further demonstrate the wider applicability of our model system for reverse engineering anatomically relevant networks for preclinical disease modelling, we used this system to culture neurons extracted from brain regions of interest from adult AD model animals. This method for the dissection and culturing of layer specific entorhinal and hippocampal neurons from AD model rats and mice builds upon our recently published protocol for extraction and culturing of LEC LII neurons from adult AD model APP/PS1 mice (39). We found that adult neurons from both AD model rats-(**Figure 5**) and mice (**Figure S2**) were able to re-form structural connections *in vitro*. Furthermore, we found that an astrocytic feeder layer was crucial for attachment of adult entorhinal and hippocampal AD neurons (**Figure S3A**). Various combinations of neural media supplements were tested (**Supplementary Table S1**), and neurites of adult rat neurons showed significantly enhanced outgrowth when adding the growth factor FGF2 (10 ng/mL) after 5 DIV (**Figure S3B** and **S3C**, all p-values for P3 < 0.01). Immunocytochemistry of the cultured adult neurons showed co-expression of neuronal marker NeuN and reelin in the LEC LII neurons, thus confirming regional identity, and co-expression of NeuN and Neurofilament Heavy for hippocampal DG-gr, CA3-pyr and CA1-pyr. This confirmed that the engineered network consisted of the neurons of interest (**Figure 5**).

**Figure 5.**
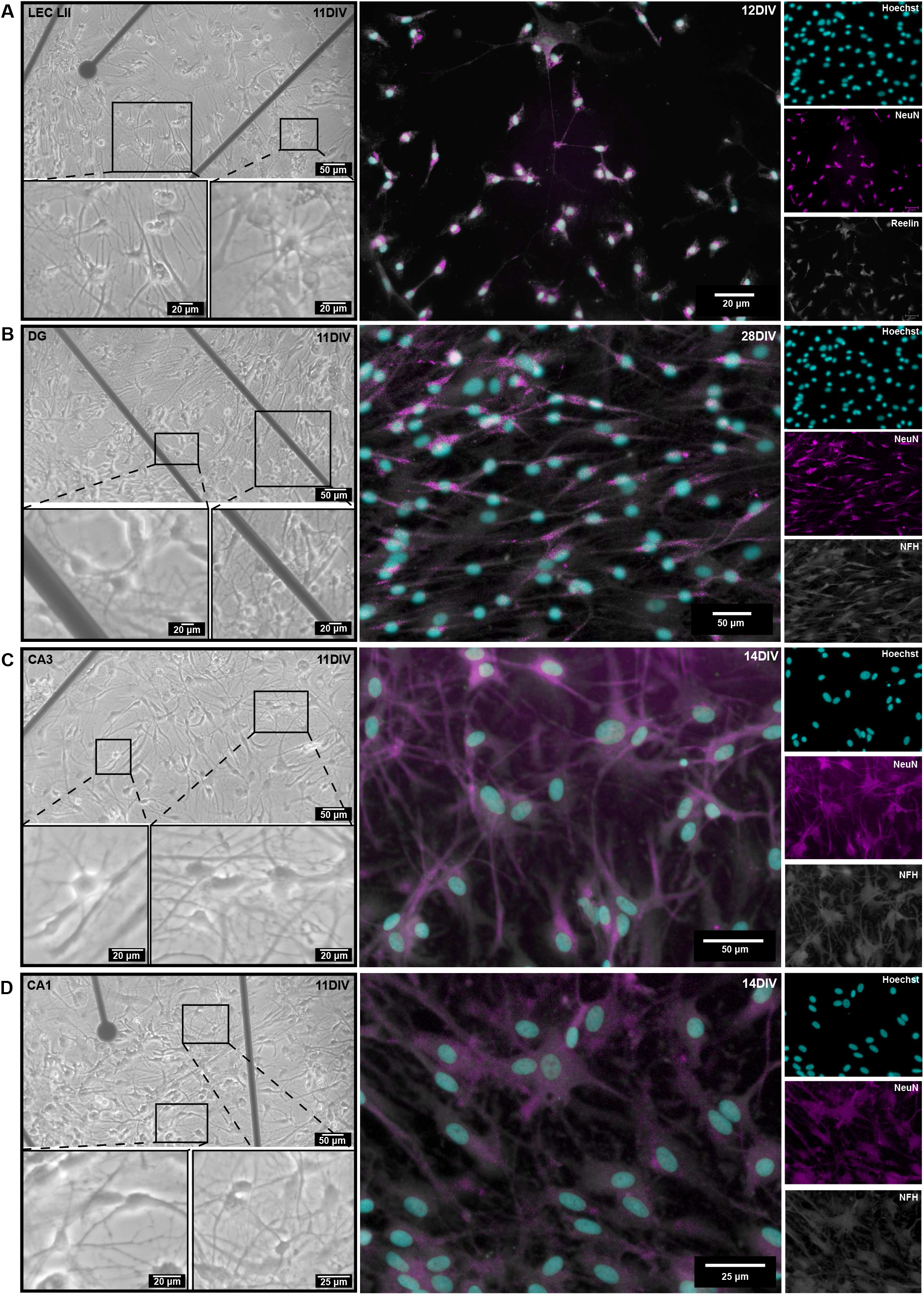
Establishment of adult entorhinal and hippocampal AD neurons on four-nodal microfluidic platforms. **(A.)** Left: Phase-contrast images of adult lateral entorhinal cortex layer II (LEC LII) neurons at 11 days *in vitro* (DIV), with enlarged insets from all regions showing neural bodies with extruding neurites. Right: Co-expression of the neural markers NeuN and reelin confirmed regional identity at 12 DIV. **(B.)** Left: Phase-contrast images of adult dentate gyrus (DG) granular neurons at 11 DIV. Right: Co-expression of neuronal markers NeuN and Neurofilament Heavy (NFH) confirmed neuronal identity at 28 DIV. **(C.)** Left: Phase-contrast images of adult Cornu Ammonis 3 (CA3) neurons at 11 DIV. Right: Co-expression of neuronal markers NeuN and Neurofilament Heavy (NFH) at 14 DIV. **(D.)** Left: Phase-contrast images of adult Cornu Ammonis 1 (CA1) neurons at 11 DIV. Right: Co-expression of neuronal markers NeuN and Neurofilament Heavy (NFH) at 14 DIV.

Adult neural networks were furthermore readily established on MEAs (**Figure 6A**). From 15 DIV, networks of both AD model rats and mice started exhibiting spontaneously evoked activity (**Figures 6B, 6C** and **S4A**, respectively). This activity was expressed as sparse, desynchronized spikes with no apparent bursting. Moreover, calculations of the median firing rate showed that neurons from both controls, homozygous and heterozygous McGill-APP rats remained active up to at least 47 days *in vitro* (**Figures 6D-6F**). Networks of neurons from AD model mice remained active up to 56 DIV (**Figure S4B**). ICC furthermore confirmed that the heterozygous and homozygous networks of neurons from McGill-APP rats expressed McSA1, indicative of A*β* aggregates (**Figure 6G**).

**Figure 6.**
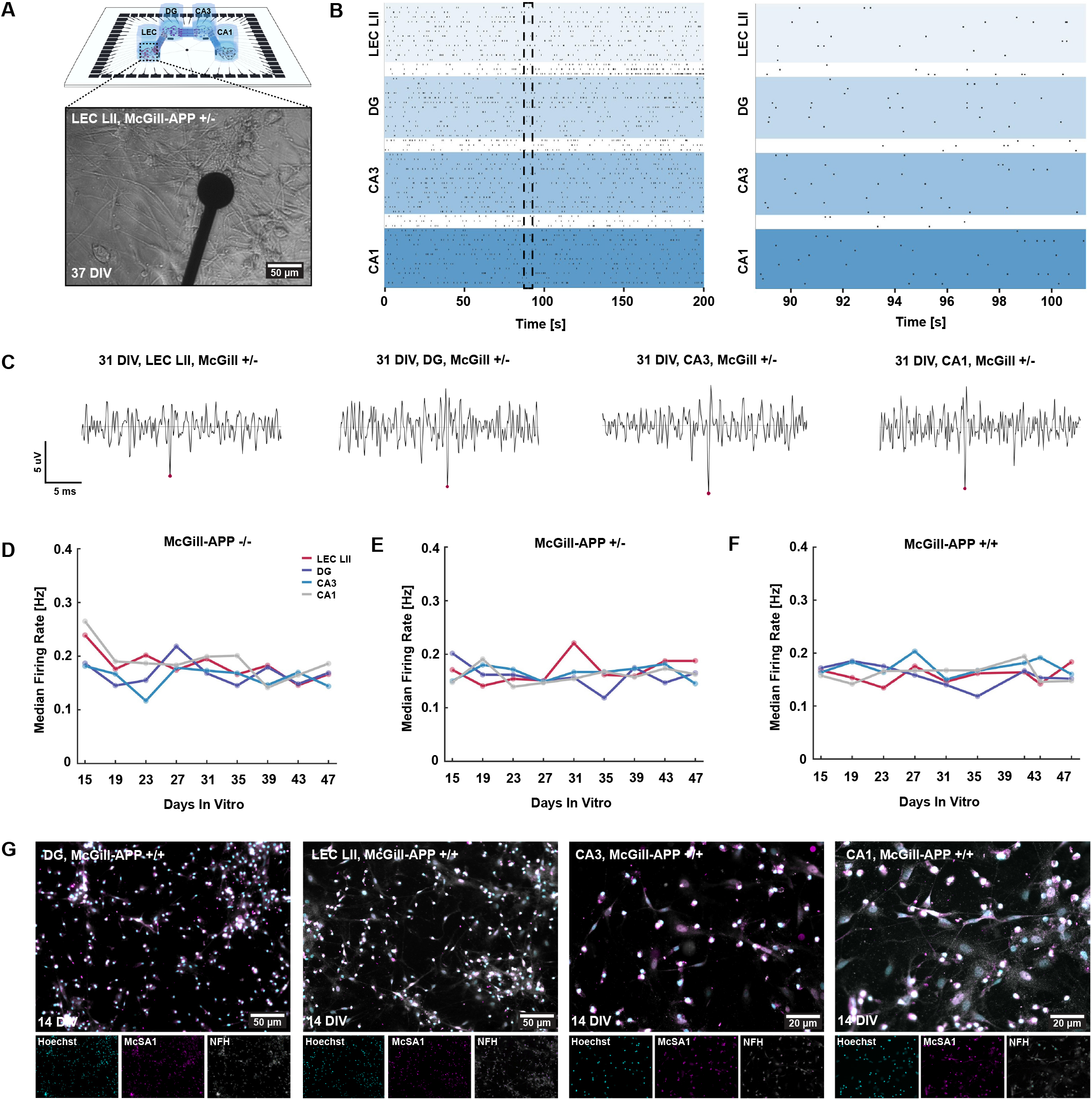
Electrophysiological characteristics of adult entorhinal and hippocampal neural networks from AD-model rats. **(A.)** Illustration of a four-nodal microfluidic chip on a microelectrode array with an inset of a phase contrast image of a lateral entorhinal cortex layer II (LEC LII) neural network derived from a McGill-APP rat +/-. **(B.)** Spike traces showing one representative spike from each subregion, LEC LII (bottom left), DG (top left), CA3 (top right) and CA1 (bottom right) at 31 DIV. **(C.)** Representative raster plots showing spontaneous network activity from all four nodes, indicating sparse, desynchronized spikes with no apparent bursting. **(D-F.)** Median firing rate (Hz) over time from 15 DIV up to 47 DIV. LEC LII; Lateral entorhinal cortex layer II. DG; Dentate gyrus. CA3; Cornu ammonis 3. CA1; Cornu ammonis 3. DIV; Days *in vitro*. McGill-APP -/-: Control rats without the APP K670M671delinsNL and APP V717F mutations. McGill-APP +/-: Heterozygous transgenic rats. McGill-APP +/+: Homozygous transgenic rats. **(G.)** Co-expression of markers for Amyloid beta (McSA1, magenta), (NFH, grey) in LEC LII and hippocampal DG, CA3 and CA1 neural networks confirm the early accumulation of A*β* in such neurons derived from an adult McGill-APP +/+ rat.

## Discussion

In this study we have demonstrated reverse engineering of complex multinodal cortical-hippocampal neural networks with controllable unidirectional connectivity. To illustrate the potential of this approach for advanced modelling of neural network function and dysfunction, we selectively induced AD relevant perturbations to healthy neural networks. Additionally, we have demonstrated establishment of layer specific entorhinal-hippocampal networks from neurons dissected from adult AD transgenic rats and mice on our advanced platforms.

For engineered neural networks to reach a mature, computationally efficient state, key design principles such as a modular microarchitecture with directional interconnectivity must be incorporated into the design of the microfluidic MEA interfaces (54–56). By incorporating Tesla valves in the microtunnels, promoting feedforward connectivity between the interconnected nodes, we showed that the reverse engineered networks organised in a computationally efficient manner. Specifically, by applying a modularity algorithm, we identified distinct communities within the individual nodes of our four-nodal networks. All networks had a high network modularity close to 0.6, which remained stable throughout the experimental period. In our previous study we found the median modularity value to be 0.39 and 0.03 for two-nodal and one-nodal networks, respectively (20). As the modularity value describes how easily neural networks can be divided into distinct communities, these results illustrate the importance of multinodal interfaces for aiding efficient network organization supporting segregated, yet functionally integrated information processing in different neuronal nodes. Our interfaces were furthermore designed to promote feedforward connectivity between the distinct communities, recapitulating directional information propagation in the brain. To evaluate the efficacy of this design in recapitulating such dynamics, we studied how network bursts initiated in the cortical, i.e. upstream, node spread downstream to the three interconnected hippocampal nodes. These bursts alternated between spanning two, three or four nodes, indicating that the networks had a capacity for gating information, i.e. selectively transmitting only a subset of the information between the distinct communities (20, 21, 27, 28). Information gating is a fundamental property of neural networks *in vivo*, conferring an ability for segregated and integrated information processing across interconnected nodes. Recapitulation of such complex network behaviour is therefore essential for modelling neural network dynamics in physiological and pathological states, including studying mechanistic effects and functional impact of neurodegenerative disease processes.

One of the major advantages of using the advanced interfaces demonstrated in this study is the ability to capture dynamical structural and functional alterations in the reverse engineered neural networks. By combining fluorescent live imaging with electrophysiology, we could longitudinally monitor the dynamic behaviour of these networks at the micro- and mesoscale. This combined approach can be challenging to implement *in vivo*. An increasing body of literature, including recent studies from our group, has characterized how *in vitro* neural networks develop and mature in healthy, i.e., unperturbed, conditions (1, 3, 20, 40, 41). Such studies provide valuable insights into the intrinsic self-organizing behaviour of engineered neural networks. In this study, we build upon these insights by reverse engineering an anatomically relevant feedforward microcircuit recapitulating key connections in the cortical-hippocampal loop. As such, our novel engineering approach renders the relevant interfaces particular suitable for a range of biomedical research applications. By selectively inducing neurodegenerative pathology associated with AD in upstream cortical nodes we could monitor network responses across the immediately affected and down-stream interconnected nodes. We show that the pathological perturbation caused increased levels of A*β* fragments and hyperactive behaviour in the cortical node, akin to what is seen *in vivo* (57). With time, developing pathology impacted network integration by reducing the information transmission between the cortical node and the downstream hippocampal nodes. These results illustrate that by reverse engineering anatomically relevant neural networks, we can model neurodegenerative pathology and its temporal progression in a controllable microenvironment.

Comparison of healthy networks to networks exhibiting inherent or induced pathological traits can furthermore be used to elucidate key disease mechanisms and therapeutic targets (58, 59). A critical risk factor for several neurodegenerative diseases and disorders, including AD, is age. While the use of embryonic cells can give fundamental insights into the impact of induced pathology, inherent pathology is better modelled using adult cells, retaining the epigenetic mature neuronal profile. By dissecting distinct cellular layers from adult animals, we could establish networks of terminally differentiated non-mitotic neurons from adult rodents with AD pathology. Specifically, we microdissected layer specific entorhinal and hippocampal neurons from adult AD model mice and rats, which are regions known to be affected during early and later stages of AD pathology. We show that the adult neural networks derived from these brain regions self-organized within a week and remained structurally connected over the span of two months. Such long-term cell viability is essential to model the gradual disease progression of neurodegeneration. Furthermore, we show that the adult neurons displayed desynchronized firing patterns with no apparent bursting. This is in line with recently reported findings by us and others from adult mouse entorhinal (39), and rat cortical and hippocampal neural networks (60) established on MEAs. Similar findings have also been reported from adult hippocampal (61, 62) and entorhinal (63) slice cultures using patch-clamp recordings. These findings show that networks established from adult neurons exhibit fundamentally different electrophysiological profiles compared to networks matured from an embryonic state *in vitro*. This needs to be taken into consideration when studying neural network behaviour in both healthy and diseased conditions. In the present study we reverse engineered entorhinal-hippocampal circuits, however, our approach is widely applicable for reverse engineering of microcircuits from other brain regions as well. Combining multiple interconnected nodes containing brain subregion- and layer specific cells enables the study of progressive disease mechanisms affecting interconnected networks, and can be used to identify potential time windows and/or modes of therapeutic intervention.

To summarize, the focus of our study was to reverse engineer multinodal neural networks with controllable afferent-efferent connectivity. Neural microcircuits in the brain, such as cortical-hippocampal networks, are profoundly complex systems with an intricate microarchitecture involving distinct subpopulations of neurons and other cells. Our approach clearly illustrates the robustness and potential of advanced neuroengineering for recapitulating and longitudinally monitoring complex network dynamics *in vitro* at the micro- and mesoscale. Furthermore, it demonstrates the power of utilizing complex network analysis approaches, such as information- and graph theory, to characterize fundamental network attributes and their importance for information processing. This enables the study of complex network dynamics in healthy and perturbed conditions, including neuropathological processes in adult neural networks. Combination with other cell types of interest thus opens for a wide range of applications in biomedical disease modelling. The potential impact of our advanced reverse engineering approach is thus manifold. It represents an integrated multidisciplinary methodology for advanced disease modelling that can accelerate elucidation of mechanistic causes of disease, pinpoint critical time points for therapeutic intervention, and support testing of the efficacy of such interventions with a view to clinical translation.

## Materials and Methods

### Experimental Design

Experimental timelines including a schematic of the microfluidic MEA design can be seen in **Figure 7**. Our neural cultures comprised either commercially available rat embryonic cortical and hippocampal neurons (hereafter referred to as cortical-hippocampal networks), or lateral entorhinal cortex layer II neurons (LEC-LII), Dentate Gyrus granular neurons (DG-gl), CA3 pyramidal neurons (CA3-pyr) and CA1 pyramidal neurons (CA1-pyr), freshly dissected from adult McGill-R-Thy1-APP AD-model rats or APP/PS1 AD-model mice (hereafter referred to as adult entorhinal-hippocampal AD networks). The method for the dissection and culturing of adult hippocampal neurons was refined from our previously reported protocol for adult lateral-most LEC LII (lLEC-LII) neurons from APP/PS1 model mice (39). For all cultures, coating and establishment of an astrocytic feeder layer was conducted using the same protocol.

**Figure 7.**
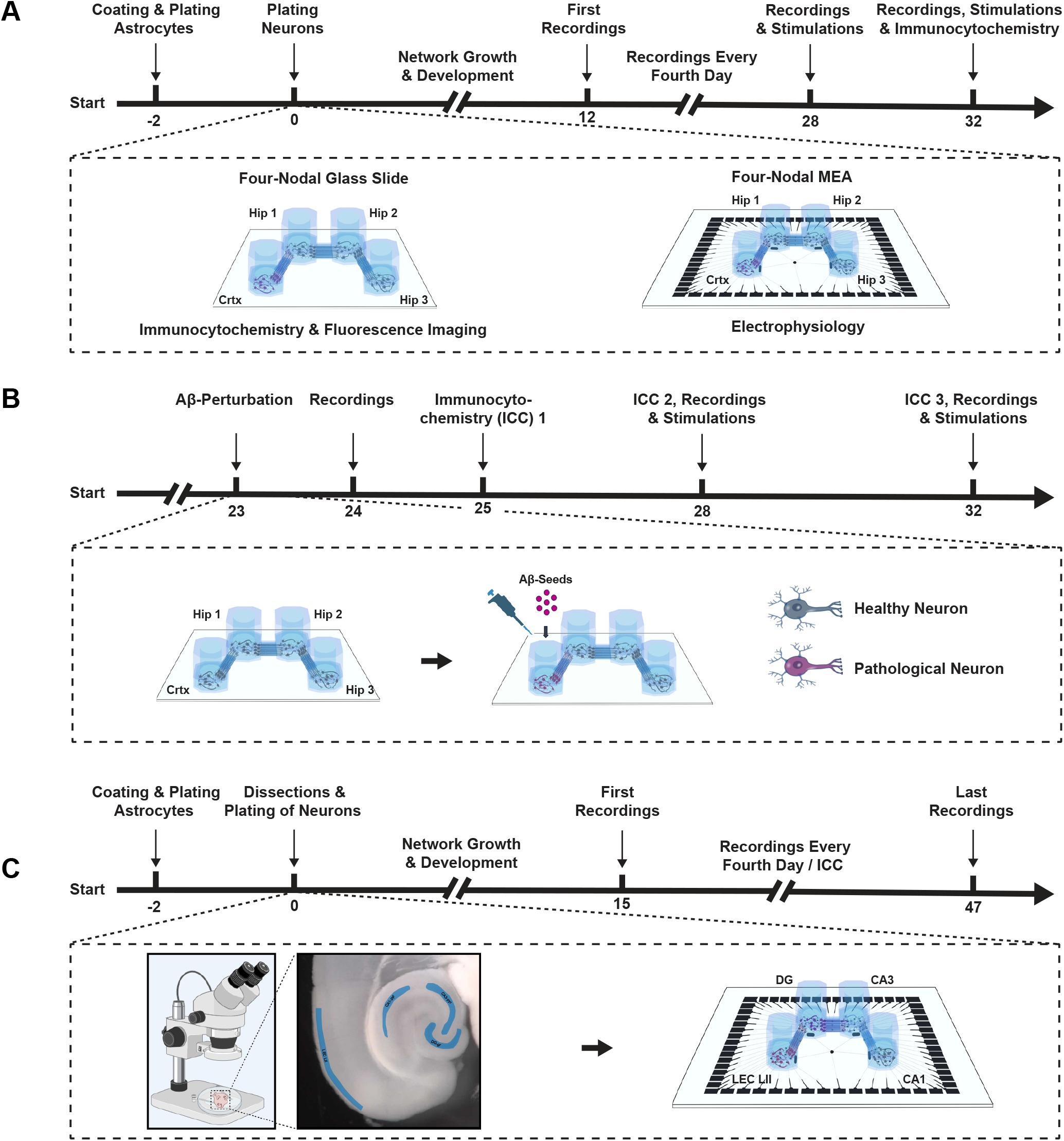
Experimental timelines. **(A.)** Four-nodal microfluidic chips were interfaced with either glass coverslips for immunocytochemistry (ICC) or microelectrode arrays (MEAs) for electrophysiology. All interfaces were coated with Poly-L-Ornithine and laminin, and a feeder layer of rat cortical astrocytes were plated two days prior to plating of commercially available rat cortical (node 1) and hippocampal (nodes 2-4) neurons. Spontaneously evoked activity was recorded every fourth day from 12 to 32 DIV, and electrical stimulations were induced after the recordings at 28 and 32 DIV. **(B.)** For perturbed networks, all steps were identical up to 23DIV. At this point, Amyloid beta (A*β*) fragments were added to the cortical node. ICC was conducted 2, 5 and 9 days after perturbation to assess the change in concentration of amyloid beta oligomers in all nodes. Recordings were furthermore conducted 1, 5 and 9 days after the perturbation to evaluate the impact on the network functionality. **(C.)** To reconstruct adult entorhinal-hippocampal circuits within the four-nodal microfluidic chips, brains of adult mice and rats were extracted and horizontally sectioned before dissection of the lateral entorhinal cortex layer II, and hippocampal subregional layers (dentate gyrus granular layer, and CA3- and CA1 pyramidal layers). Dissected entorhinal and hippocampal tissues were next dissociated into single cells enzymatically and manually by trituration. LEC LII, DG-gl, CA3-pyr and CA1-pyr were plated in nodes 1-4, respectively. A horizontally cut brain section viewed under a stereoscope using fiber optic illumination for contrast with delineation of entorhinal and hippocampal subregions is illustrated below the timeline. Lateral entorhinal cortex layer II (LEC LII) and hippocampal dentate gyrus granular layer (DG-gl), CA3 pyramidal layer (CA3-pyr) and CA1 pyramidal layer (CA1-pyr) are highlighted in blue.

### Design & Fabrication of Microdevices

The four-nodal microdevices were designed using Clewin 4 (WieWeb Software, Enschede). Each node was 5 mm in diameter and 60 µm high. Furthermore, the nodes were connected by 20 microtunnels. To induce unidirectional axonal outgrowth between the nodes, a geometrical design inspired by the Tesla valve was implemented within the microtunnel architecture (20). All tunnels were 350 µm long, 10 µm wide and 5 µm high. The design also had spine structures on the postsynaptic side to misguide axons trying to enter the microtunnel inlets. 50 nanoporous platinum electrodes of 50 µm diameter were positioned evenly spread across the four nodes. Additionally, 9 electrodes were positioned within the microfluidic tunnels to confirm active connections between the nodes. A reference electrode was also positioned in each of the four nodes, and all four reference electrodes were connected to the same tunnel outlet on the Multichannel system. The design, as well as an image of a fabricated microdevice can be seen in **Figure S5**. Fabrication of all microdevices was conducted according to our recently reported protocol (20).

### Coating of Culturing Platforms

Prior to coating, all microdevices were sterilized in UV light overnight in a biosafety cabinet. Consecutively, the distilled (DI) water in the devices was replaced by Poly-L-Ornithine solution (PLO) (Sigma-Aldrich, A-004-C) at a concentration of 0.1 mg*/*mL and incubated at 37 *°*C, 5 % CO_2_ for 2 h *or* overnight at 4 *°*C. Subsequently, all PLO was discarded and the microdevices rinsed three times for 10 min with milli-Q (MQ) water. After the last rinse, laminin solution consisting of 16 µg*/*mL natural mouse laminin (Gibco™, 23017015) in phosphate-buffered saline (PBS, Sigma-Aldrich, D8537) was added and the microdevices incubated at 37 *°*C, 5 % CO_2_ for 2 h. A hydrostatic pressure gradient was established during all coating to ensure flow of the coating solution through the microtunnels. This was achieved by filling the chambers with an unequal amount of coating solution.

### Seeding of Astrocyte Feeder Layer

For the astrocytic feeder layer, a solution consisting of DMEM, low glucose (Gibco™, 11885084) supplemented with 15 % Fetal Bovine Serum (Sigma-Aldrich, F9665) and 2 % Penicillin-Streptomycin (Sigma-Aldrich, P4333) was prepared. Next, the coating solution was replaced by the astrocyte media, and rat cortical astrocytes (Gibco™, N7745100) were seeded at a density of 100 cells*/*mm2, i.e., 2000 cells per microchamber. The astrocytes were allowed to expand for 48 h, before seeding of either embryonic cortical and hippocampal neurons or adult entorhinal and hippocampal neurons.

### Embryonic Neurons: Plating and Maintenance

Neural media consisting of Neurobasal Plus Medium (Gibco™, A3582801) supplemented with 2 % B27+ (Gibco™, A358201), 1 % GlutaMAX (Gibco™, 35050038) and 2 % Penicillin-Streptomycin (Sigma-Aldrich, P4333) was prepared. Rock Inhibitor (Y-27632 dihydrochloride, Y0503, Sigma-Aldrich) was additionally added to the media during the first two days of neural growth at a concentration of 0.1 % to increase neural survival. Rat cortical neurons from Sprague Dawley rats (Gibco, A36511) were plated at a density of 1000 cells*/*mm2, equaling 20 000 cells in the first node of each microfluidic interface. The cortical neurons were thereafter allowed to settle in an incubator for 3 h before plating the hippocampal neurons. Next, rat hippocampal neurons from Sprague Dawley rats (Gibco, A36513) were plated at a density of 1000 cells*/*mm2 in the three remaining nodes of each microfluidic interface. Half the neural media was replaced with fresh neural media 4 h after plating, and again after 24 h. From here on, half the neural media was replaced every second day throughout the experimental period.

### Viral Transductions

Viral transductions for inducing ubiquitous expression of fluorescent markers were used to verify unidirectional structural connectivity between the nodes, according to our previously reported method (20). Viral vectors were prepared in-house at the Viral Vector Core Facility, NTNU. At 11 DIV, neural networks (n = 4) were transduced with AAV 2/1 serotype viruses loaded with either pAAV-CMV-beta Globin intron-EGFP-WPRE-PolyA or pAAV-CMV-beta Globin intron-mCherry-WPRE-PolyA under a CMV promoter to ubiquitously express GFP (nodes 1 and 3) or mCherry (nodes 2 and 4). 3/4 of the media in nodes 1 and 3 were removed to create a hydrostatic pressure gradient across the nodes, and viruses for expression of GFP were added at a concentration of 5*e*^2^ viruses/cell (i.e. 1*e*^7^ viruses per node). The cells were subsequently incubated for 3 h at 37 *°*C and 5 % CO_2_. Next, nodes 1 and 3 were filled all the way up with cell media, and 3/4 of the media in nodes 2 and 4 was removed before adding viruses to express mCherry at a concentration of 5*e*^2^ viruses/cell. After 3 h of incubation, these nodes were also filled all the way up with media. Imaging of the networks were conducted at 28 DIV.

### Amyloid Beta Perturbations

At 23 DIV, the cortical node in six four-nodal networks were perturbed using A*β*-fragments. 1 mg Beta-Amyloid Peptide (1-42, human) (ab120301, abcam) was dissolved in 100 µL 1 % NH_4_OH (458680025, Thermo Scientific). The solution was furthermore diluted to 2 mM in neuronal media. This concentration was chosen based on previous literature (64), and initial concentration testing (**Figures S6** and **S7**), to induce A*β*-oligomerization while still retaining network viability over prolonged periods of time. Subsequently, half the cell media in the cortical node was replaced by 2 mM A*β*-solution. Immunocytochemistry was conducted 2, 5 and 9 days after the perturbations to assess the change in concentration of A*β*-deposits in all nodes.

### Animals Models and Genotyping

All animal experiments comply with the ARRIVE guidelines (65), were approved by the Norwegian Food Safety Authority and carried out in accordance with the EU (European Union) Directive 2010/63/EU for animal experiments. The animals were held in standard lab cages (up to 5 animals per cage), with temperatures of 22 ± 2 *°*C, and kept at a light/dark cycle of 12:12 hours with access to food and water *ad libitum*. For optimization of the protocol we used a total of 43 adult McGill-R-Thy1-APP rats, either homozygote (+/+), heterozygote (+/-) or negative genotype (-/-), including 22 female and 21 male. Of these, we collected data from 14 rats (8 female and 6 male) and from a total of 2 female, adult APP/PS1 mice. See **Supplementary Table S2** for a total overview of sex, age and genotype of all animals.

The APP/PS1 mouse model is a transgenic familial AD-model, with a C57BL/6J genetic background, co-expressing mutated human amyloid precursor protein (hAPP) (KM670/671NL) and mutated human presenilin 1 (L166). Both mutated proteins are driven by expression cassettes under the Thy1 promoter. This causes increased levels of Aβ leading to formation of extracellular cortical Aβ-deposits starting already from around 6 weeks of age (66). The APP/PS1 mice were genotyped using a KAPA-kit as described in Hanssen *et al*. (39). The McGill-R-Thy1-APP rat model is a transgenic familial AD model with a Wistar (HsdBrl:WH) genetic background, co-expressing mutated hAPP (APP K670M671delinsNL) and (APP V717F) (67).

The McGill-R-Thy1-APP rats were genotyped using quantitative PCR (qPCR) and genomic DNA isolated from ear tissue as described in previous studies (68). Both homozygous and heterozygous rats were included in the study, in addition to controls.

### Adult Neurons: Dissection

All tools were sterilized by autoclaving before use. Both mice and rats were deeply anesthetized using 5 % isoflurane gas (Abbott Lab, 05260–05) and checked for absence of reflexes before the head was decapitated with a guillotine for rats or surgical scissors (FST, 14007–14) for mice. Subsequently, dissection and sectioning of the brain were conducted as described previously for both mice and rats. See Hanssen *et al*. for a detailed description of the procedure (39). Brains were extracted in a petri dish filled with Hanks Balanced Salt Solution (Thermo Fischer Scientific, 88284) kept on ice. Extracted brains were sectioned horizontally on a Leica VT1000 S vibratome (Leica Biosystems) (thickness of 300 µm, frequency of 5.3 Hz and speed of 8 mm*/*s) with 0.2 % AGAR (VWR, 20767.298) supporting the caudal end of the brain. Sectioned brain slices were immediately transferred to wells of a sterile 24-well plate held on ice. Sections were further transferred to a petri dish held on ice and viewed under a stereoscope with fiber optic lamps, allowing for sufficient contrast of the cytoarchitectonic landmarks. Dissection of the lateral entorhinal cortex layer II (LEC-LII) was conducted as described in our recent study for both mice and rats (39). Additionally, the dentate gyrus granular layer (DG-gl), CA3 pyramidal (pyr) and CA1-pyr layers were dissected (**Figure 7C**). The DG-gl can be separated from the molecular layer and hilus using its densely packed layer of granular neurons. Similarly, the pyramidal layer in CA3 and CA1 consists of densely packed pyramidal neurons that can be distinguished from superficial layers lucidum, radiatum and lacunosum-moleculare and deep layer oriens (29). All dissected strips of tissue were placed by region in four separate sterile 15 mL tubes filled with dissection media (HABG-P) consisting of Hibernate-A (Thermo Fischer Scientific, A1247501) supplemented with 2 % B27+ (Gibco™, A3582801), 2.5 % GlutaMAX (Gibco™, 35050061) and 1 % Penicillin-streptomycin (Sigma-Aldrich, P4333), placed on ice.

### Adult Neurons: Plating and Maintenance

All dissected tissue was rinsed for debris three times by adding and removing 3 mL of HABG-P. Following this, all media was discarded and the tissue was dissociated for 15 min at 37 *°*C, 5 % CO_2_, 20 % O_2_ in Neural Isolation Enzyme Papain (Thermo Fischer Scientific, 88285) reconstituted in Hank’s Balanced Salt solution (Thermo Fischer Scientific, 88284) as described by the manufacturer (Thermo Fischer Scientific, MAN0011896). Subsequently, all dissociation enzyme was discarded, 100 µL HABG-P was added and the tissue was manually triturated 10 times by using a 100 µL pipette tip. Further, HABG-P was topped up to a total of 3 mL and the vials of tissue were centrifuged at x200 g for 2 min in room temperature. After centrifugation, HABG-P was discarded and 100 µL neural media was added before manual trituration 10 times using a 100 µL pipette tip. The neural media consisted of Neurobasal Plus Medium (Gibco™, A3582801) supplemented with 2 % B27+ (Gibco™, A358201), 1 % GlutaMAX (Gibco™, 35050038), 2 % Penicillin-Streptomycin (Sigma-Aldrich, P4333), 0.1 % Rock Inhibitor (Y-27632 dihydrochloride, Y0503, Sigma-Aldrich) and 10 % Fetal Bovine Serum (Sigma Aldrich, 12106C). For adult neurons derived from mice, 0.1 % BDNF (Neurotrophins, 450-02) was added to the neural media. For adult neurons derived from rats, 0.1 % FGF2 (Thermo Fisher Scientific, 13256-029) was added to the neural media. Neurons were plated at a density of 750 cells*/*mm2, equaling 15 000 cells in each node of the microfluidic chip. Half the neural media was replaced every second day throughout the experimental period.

### Immunocytochemistry

For immunocytochemistry, both cortical-hippocampal and adult entorhinal-hippocampal neurons were plated in microfluidic platforms bonded to glass coverslips (VWR International, 24x24 mm No. 1 Menzel-Gläser). Fixation of cortical-hippocampal cultures was conducted using glyoxal solution based on the protocol by Richter *et al*. (69). This fixative consisted of 20 % ethanol absolute (Kemetyl, 100 %), 8.0 % Glyoxal solution (Sigma-Aldrich, 128465) and 1 % acetic acid (Sigma-Aldrich, 1.00063) in MQ-water. Fixation of adult entorhinal-hippocampal cultures was conducted using PBS (Sigma-Aldrich, D8537) containing 0.4 M phosphate buffer and 4 % freshly depolymerized paraformaldehyde (PFA; Sigma-Aldrich, P6148). The cultures were washed with PBS to remove debris, before the fixative was applied for 15 min at room temperature. Subsequently, all chambers were washed with PBS 3 times for 15 min each. Next, 0.5 % Triton-X (Sigma-Aldrich, 1086431000) diluted in PBS was applied to permeabilize the cells. All chambers were again washed twice with PBS before a blocking solution consisting of 5 % goat serum (Abcam, ab7481) diluted in PBS was added to the cultures, and the cultures incubated at room temperature on a shaking table at 30 rpm for 1 h. Primary antibody solutions were prepared in PBS with 5 % goat serum, and antibody concentrations according to the ones listed in **Table** Cultures were placed on a shaker table at 30 rpm at 4 *°*C overnight. Following this, the primary antibody solution was removed, and the cell cultures were rinsed three times with PBS for 15 min each. Next, a secondary antibody solution consisting of 0.2 % secondaries and 5 % goat serum diluted in PBS was added to the cultures. The cultures were left to rest on a shaker table at 30 rpm for 3 h at room temperature. Prior to applying the secondary antibody solution, the prepared solutions were centrifuged at 6000 rpm for at least 15 min to remove precipitates. Subsequently, the secondary antibody solution was replaced by Hoechst (Abcam, ab228550) at a concentration of 0.1 % diluted in PBS, and the cultures left on a shaker table for another 30 min. Eventually, all cultures were washed three more times in PBS, then twice in MQ water prior to imaging. All images were acquired using an EVOS M5000 microscope (Invitrogen). The microscope was equipped with DAPI (AMEP4650), CY5 (AMEP4656), GFP (AMEP4651) and TxRed (AMEP4655) LED light cubes and Olympus UPLSAP0 4X/0.16 NA and 20x/0.75 NA objectives. Post-processing of images was conducted in ImageJ/Fiji or Adobe Photoshop 2020.

**Table 1.** Antibodies and concentrations used for Immunocytochemistry.

| Marker | Catalogue Number | Concentration |
| --- | --- | --- |
| β3-Tubulin | Ab78078 | 1/1000 |
| GAP-43 | 9264 (Sigma-Aldrich) | 1/500 |
| GFAP | Ab7260 | 1/1000 |
| MAP2 | Ab5392 | 1/5000 |
| McSA1 | MM-0015-P (MédiMabs) | 1/750 |
| NeuN | Ab279295 | 1/500 |
| Neurofilament Heavy | Ab4680 | 1/5000 |
| Reelin | Ab78540 | 1/10000 |

All antibodies were purchased from Abcam unless otherwise stated.

### Quantification of Amyloid Beta Deposits

For quantification of amyloid beta deposits, ImageJ/Fiji was utilized. All images were converted to 8-bit and thresholded using maximum entropy. Furthermore, the images were binarized, and particles separated using the watershed function. The built-in particle analysis function was subsequently used to count the number and size of the A*β*-seeds, and only particles above 5 µm were included in the final analysis. The threshold was set this high to assure only aggregates of amyloid beta were included in the analysis, and not noise caused by antibody precipitates or autofluorescence. This provided the best comparison of amyloid beta levels between networks stained at different time points.

### Imaging and Quantification of Neurite Length

For quantification of neurite length, phase contrast images acquired using a Zeiss Axio Vert V.1 brightfield 20x/53 WD/0.4 NA objective with an Axiocam 202 mono were used. For analysis, Neurolucida (MFB Bioscience) was used to quantify neurite length of the adult neurons.

### Electrophysiological Recordings

A MEA2100 workstation (Multichannel Systems) was utilized for all recordings with a sampling rate of 25000 Hz. An external temperature controller (TC01, Multichannel Systems) was used to maintain the temperature at 37 *°*C. All cultures were allowed to equilibrate at the workstation for 5 min prior to recordings, and recordings lasted for 15 min for cortical-hippocampal networks and 10 min for adult entorhinal-hippocampal neural networks. A 3D-printed plastic cover with a gas-permeable membrane was used to keep the neural networks in a sterile environment during the recordings.

### Electrical Stimulations

Stimulations of cortical-hippocampal neurons were performed following the recordings at 28 and 32 DIV using a train of 60 spikes at *±* 800 mV (positive phase first) of 200 µs duration with an interspike interval of 5 s. The most active electrode measured in spikes per second in the cortical cell population was chosen for stimulations. This was done to assure that the electrode chosen for stimulation had sufficient coupling with a neuron to induce activity.

### Data Analysis

All data preprocessing and analysis was conducted in Matlab R2021b, and graphs were plotted using the matlab function linspecer (70), based on the color palette colorBrewer (71). A 4th order Butterworth bandpass filter was used to remove noise below 300 Hz and above 3000 Hz, and a notch filter to remove noise at 50 Hz from the power supply mains. All filters were run using zero-phase digital filtering. For spike detection, the Precise Timing Spike Detection (PTSD) algorithm developed by Maccione *et al*. (72) was utilized. A threshold of 9 times the standard deviation was chosen for the cortical-hippocampal cultures, with a maximum peak duration of 1 ms and a refractory period of 1.6 ms. For the adult entorhinal-hippocampal cultures, a standard deviation of 7.5 was chosen due to the lower signal-to-noise ratio of the activity in these cultures overall. The *SpikeRasterPlot* function developed by Kraus (73) was adapted and used for creating raster plots.

Burst detection was conducted using the logISI algorithm developed by Pasquale *et al*. (40). A minimum of four consecutive spikes was set as a hard threshold for a burst to be detected, and the maximum interspike interval to 100 ms. Network bursts were detected using the logIBEI approach (74), with a minimum of 10 % of all active electrodes required to exhibit bursting activity for a network burst to be classified. After binning the data into 50 ms time bins, functional connectivity was analysed using Pearson correlation. Intranodal correlation was evaluated by taking the average correlation of all active connections within each node. Internodal correlation was evaluated by taking the correlation between the summed activity of all four nodes in the four-nodal network. To visualize key network features, graphs were plotted using the electrodes as nodes, and pairwise mutual information as the edges (i.e. lines connecting the individual nodes). The pairwise mutual information was calculated using the information theory toolbox developed by Timme & Lapish (75). Edges with weaker connectivity than 0.1 were removed from the graph representations to highlight only the strongest, direct connections. Firing rate (Hz) was furthermore used to represent the node color and pagerank centrality the node size. The Louvain algorithm was used to perform community detection, delineating the nodes into distinct communities based on which nodes were more highly interconnected with each other than the rest of the network (76). The node edge color of the graphs were furthermore used to showcase which nodes belonged to the same community. The modularity value was used to assess how easily the algorithm managed to classify the nodes into distinct communities. All graph theoretical measures were calculated using the Brain Connectivity Toolbox developed by Rubinov & Sporns (77). To remove stimulation artifacts, stimulation data was run through the SALPA filter developed by Wagenaar *et al*. (78). Additionally, 15 ms of the filtered data was blanked following each stimulation time point. Subsequently, the data was binned into 20 ms time bins, and peristimulus time histograms of the average response of the stimulations in each node plotted.

### Statistical Analysis

Generalized Linear Mixed-Effect Models (GLMMs) were used to assess differences in network parameters between control networks and networks perturbed with A*β*-fragments. The GLMMs were generated using SPSS version 29.0.0.0. The network type, i.e. perturbed networks from DIV 24 to 32 (i.e. after perturbation) or control networks (i.e. all other data points for both perturbed and unperturbed networks) were used as a fixed effect. The network age was furthermore used as a random effect, and the network feature of interest as a target. To fit the data, a gamma distribution with a log link function was chosen based on the Akaike information criterion. Sequential Bonferroni adjustment was selected for multiple comparison. All other statistical analyses were conducted using a two-sided Wilcoxon rank sum test with the Matlab function *ranksum*.

## AUTHOR CONTRIBUTIONS

The author contributions follow the [CRediT](https://credit.niso.org/) system. **KSH**: Conceptualization, Methodology, Investigation (dissections, cell experiments, electrophysiology, formal analysis), Writing – Original Draft, Visualization. **NWH**: Conceptualization, Methodology, Software, Investigation (chip design & manufacturing, cell experiments, electrophysiology, formal analysis), Writing – Original Draft, Visualization. **SNN**: Investigation (dissections, cell experiments, electrophysiology), Writing – Review & Editing. **AKF**: Writing - Review Editing, Supervision. **MPW**: Conceptualization, Funding acquisition, Writing - Review Editing, Supervision. **IS, AS**: Conceptualization, Methodology, Writing – Review & Editing, Funding acquisition, Supervision.

## ACKNOWLEDGEMENTS

We would like to thank Hanne Mali Møllegård for breeding of all APP/PS1 mice and Grethe Mari Olsen for breeding of all McGill-R-Thy1-APP rats. NTNU Enabling technologies, The Liaison Committee for education, research and innovation in Central Norway, the Olav Thon Foundation, Samarbeidsorganet HMN-NTNU (23166), K.G. Jebsen Center for Alzheimer’s Disease, and Civitan Norwegian Research Fund for Alzheimer’s Disease are acknowledged for funding this research. The Research Council of Norway is acknowledged for the support to the Norwegian Micro- and Nano-Fabrication Facility, NorFab, project number 295864. Prof. Michela Chiappalone and Prof. Sergio Martinoia, University of Genova for generously providing the scripts for the Precise Timing Spike Detection algorithm and the logISI burst detection. We would also like to thank Dr. Rajeevkumar Nair Raveendran at the Viral Vector Core Facility, Kavli Institute for systems neuroscience, for designing and preparing the AAV2/1 viruses used for the structural analysis.

## COMPETING FINANCIAL INTERESTS

The authors declare that the research was conducted in the absence of any commercial or financial relationships that could be construed as a potential conflict of interest.

## Notes

### Competing Interest Statement

The authors have declared no competing interest.

### Summary of Updates

All figures updated with extended data to strengthen the conclusions of the manuscript. Supplementary files updated.

