## Supplementary File for "Reverse Engineering of Feedforward Cortical-Hippocampal Microcircuits for Modelling Neural Network Function and Dysfunction"

### Supplementary Materials

**Katrine Sjaastad Hanssen*[1,2,✉], Nicolai Winter-Hjelm*[1,✉], Salome Nora Niethammer[1,2], Asgeir Kobro-Flatmoen[2,3], Menno P. Witter[2,3], Axel Sandvig[1,4], and Ioanna Sandvig[1,✉]**

[1]Department of Neuromedicine and Movement Science, Faculty of Medicine and Health Sciences, Norwegian University of Science and Technology (NTNU), Norway
[2]Kavli Institute for Systems Neuroscience, Centre for Neural Computation, Egil and Pauline Braathen and Fred Kavli Centre for Cortical Microcircuits, NTNU, Norway
[3]K.G. Jebsen Centre for Alzheimer's Disease, Faculty of Medicine and Health Sciences, NTNU, Norway
[4]Department of Neurology and Clinical Neurophysiology, St Olav´s University Hospital, Trondheim, Norway

**Table 1** | Overview of media supplement compositions in different protocols used throughout the study.

| Protocol 1 | Protocol 2 | Protocol 3 | Protocol 4 | Protocol 5 | Protocol 6 |
|---|---|---|---|---|---|
| NPM | NPM | NPM | NPM | NPM | NPM |
| 0.2 % B27+ | 0.2 % B27+ | 0.2 % B27+ | 0.2 % B27+ | 0.2 % B27+ | 0.2 % B27+ |
| 0.025 GlutaMAX | 0.025 GlutaMAX | 0.025 GlutaMAX | 0.025 GlutaMAX | 0.025 GlutaMAX | 0.025 GlutaMAX |
| 0.001 % BDNF | 5 ng/mL FGF2 | 10 ng/mL FGF2 | 10 ng/mL FGF2 | 0.001 % BDNF | 10 ng/mL FGF2 |
| 0.01 % Penstrep | 0.01 % Penstrep | 0.01 % Penstrep | 10 ug/mL Gentamycin | 0.01 % Penstrep | 0.01 % Penstrep |
| 0.001 % RI | 0.001 % RI | 0.001 % RI | 0.00 % RI | 0.001 % RI | 0.001 % RI |
| 0.1 % FBS | 0.1 % FBS | 0.1 % FBS | 0.0 % FBS | 0.1 % FBS | 0.1 % FBS |

*NPM; Neurobasal Plus Medium. B27+; B27 Plus Supplement. BDNF; Brain Derived Neurotrophic Factor. FGF2; F Growth Factor 2. PS; Penicillin-Streptomycin. RI; Rock Inhibitor. FBS; Fetal Bovine Serum.*

**Table 2** | Overview of animals in the study.

| Animal model | Number | Sex | Age | Genotype |
|---|---|---|---|---|
| McGill-R-Thy1-APP rat | 5 | M | P38-P89 | Heterozygote +/- |
| McGill-R-Thy1-APP rat | 4 | M | P46-P53 | Homozygote -/- |
| McGill-R-Thy1-APP rat | 12 | M | P43-P78 | Homozygote +/+ |
| McGill-R-Thy1-APP rat | 14 | F | P39-P81 | Heterozygote +/- |
| McGill-R-Thy1-APP rat | 1 | F | P53 | Homozygote -/- |
| McGill-R-Thy1-APP rat | 7 | F | P37-P72 | Homozygote +/+ |
| APP/PS1 mouse | 2 | F | P79-P80 | APP/PS1 +/- |

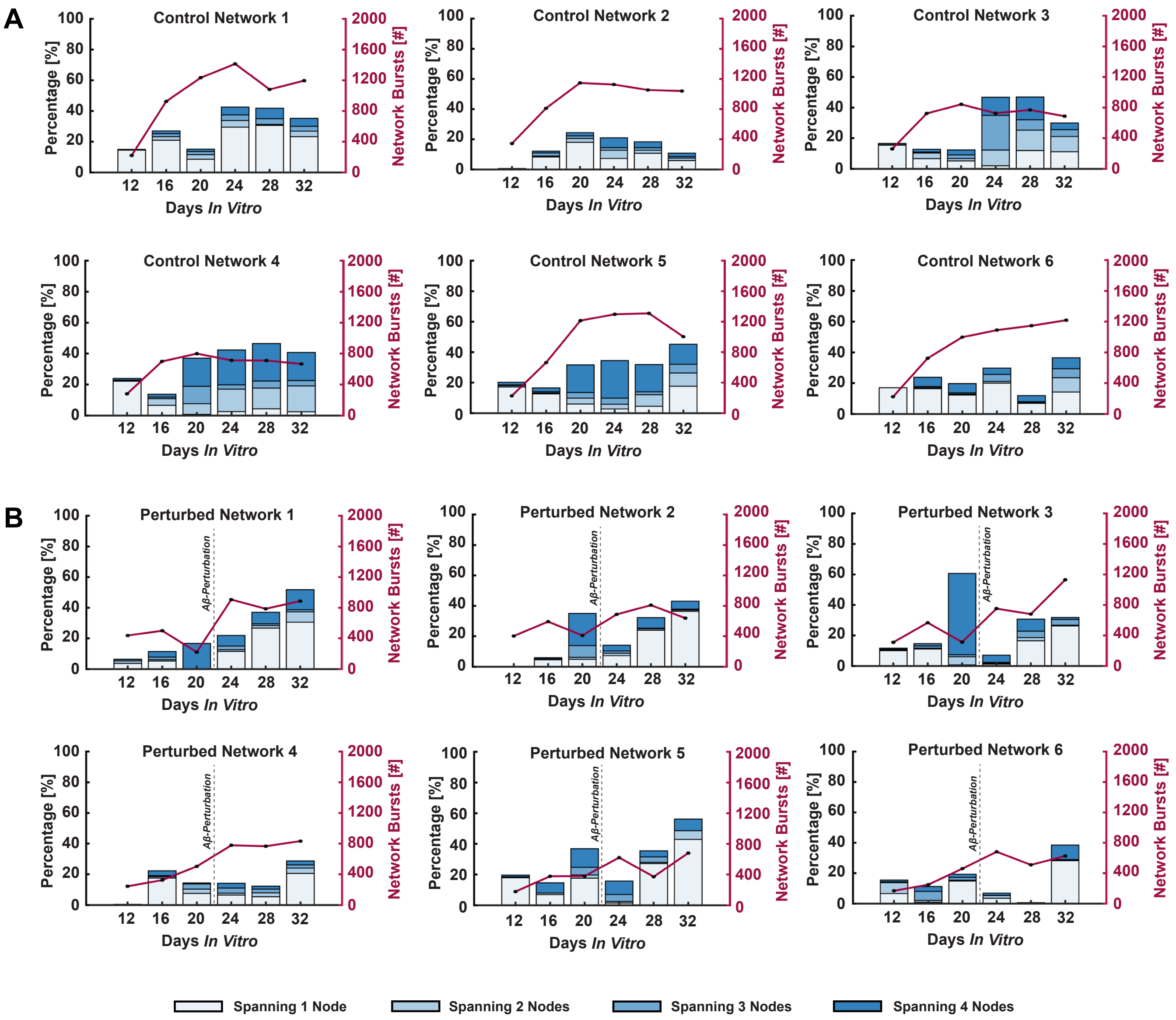


**Figure S1** | **Spread of network bursts initiated in the cortical node through the hippocampal nodes. (A.)** Histograms showing the percentage of spontaneously evoked network bursts initiated within the cortical population, and the number of nodes they span for the control networks. Each subfigure represents the activity from a single MEA. The total number of network bursts initiated during a $15\,\mathrm{min}$ recording at each DIV is shown along the secondary y-axis. **(B.)** Histograms showing the percentage of spontaneously evoked network bursts initiated within the cortical population, and the number of nodes they span for the networks perturbed with A$\beta$-fragments at 23 DIV.

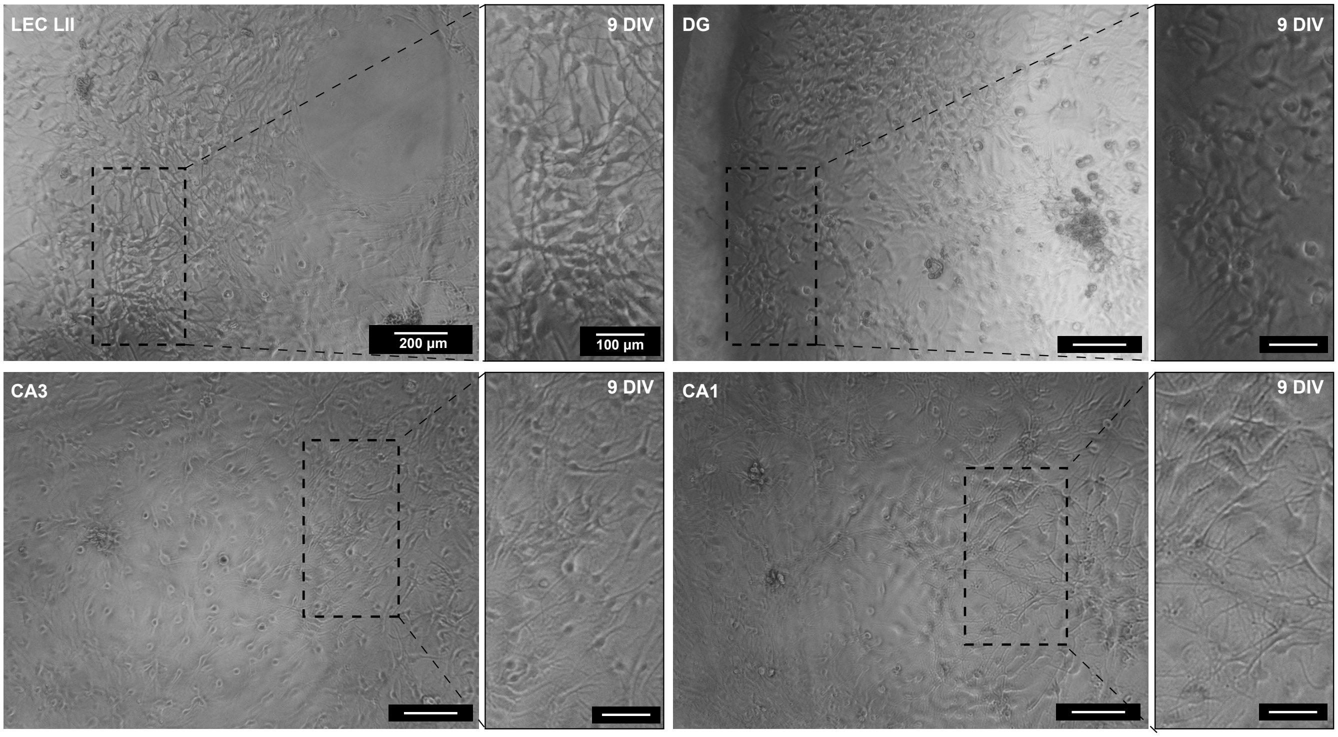


**Figure S2** | **Adult mouse neurons re-form structural connections *in vitro*.** Phase-contrast images of adult neurons from the APP/PS1 mouse model at 9 DIV. LEC LII; lateral entorhinal cortex layer II. DG; dentate gyrus. CA3/CA1; Cornu Ammonis 3/1. DIV; *days in vitro*.

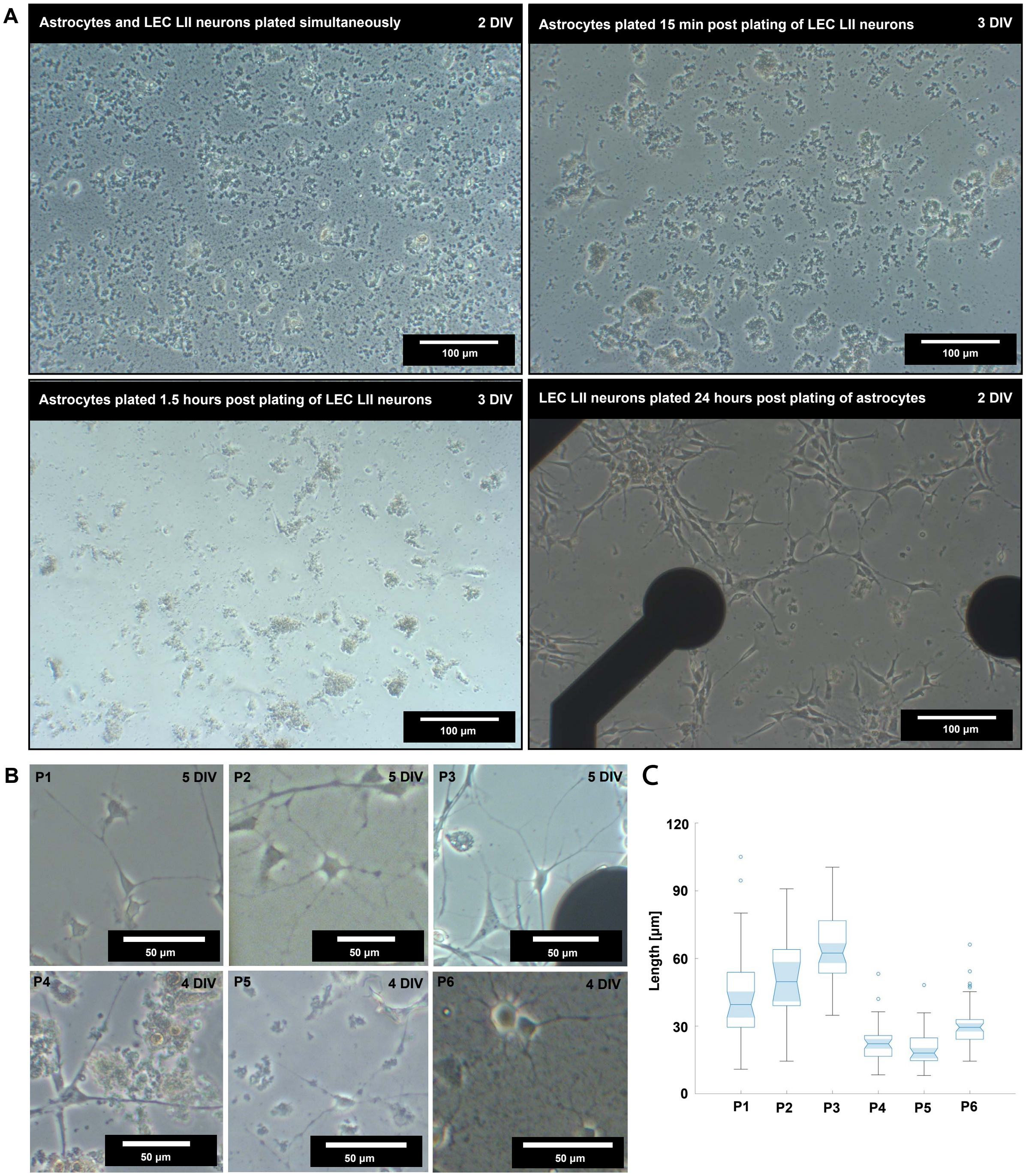


**Figure S3** | **Importance of an astrocytic monolayer and media supplement combination for viability of adult neurons from transgenic AD model rats. (A.)** Phase contrast images of astrocytes and LEC LII neurons plated simultaneously (top left), LEC LII neurons plated $15\,\mathrm{min}$ prior to plating astrocytes (top right), LEC LII neurons plated $1.5\,\mathrm{h}$ prior to plating astrocytes (bottom left) and astrocytes plated $24\,\mathrm{h}$ prior to plating LEC LII neurons (bottom right). **(B.)** Phase contrast images of cultured neurons on six different media supplement protocols, exhibiting clear differences in the length of outgrowing neurites. **(C.)** Bar graph showing the mean neurite length for each culture in **(B.)** at 4 and 5 days *in vitro*. A total of 285 neurites were measured and analyzed using Neuroucida. P1-6: Protocol 1-6.

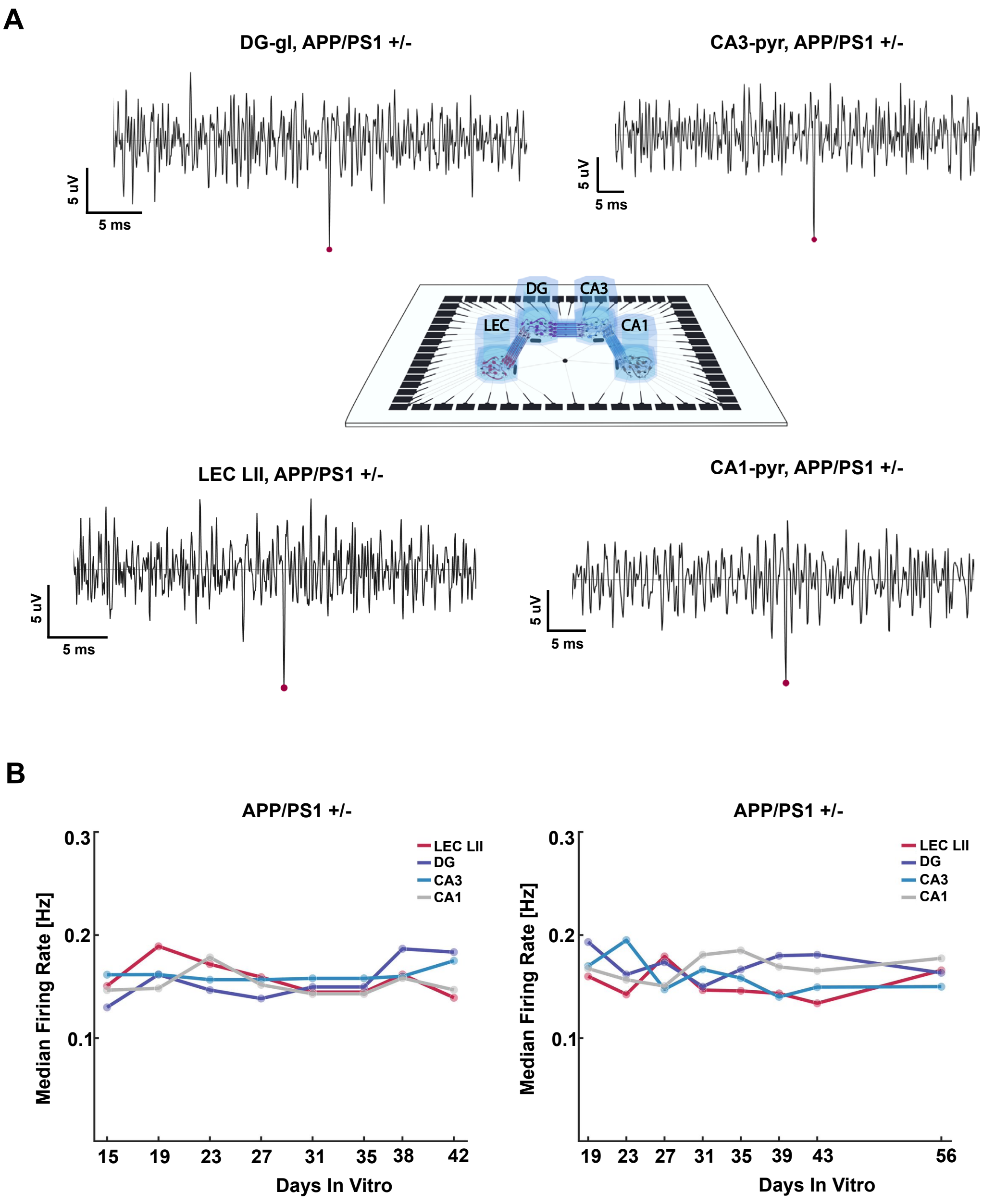


**Figure S4** | **Adult neurons from transgenic APP/PS1 mice exhibit electrophysiological activity *in vitro*. (A.)** Illustration of the four-nodal microfluidic microelectrode array (mMEA) with spike traces showing one detected spike from each entorhinal and hippocampal subregion. **(B.)** Graph of median firing rate (spikes/second) from 15 to 42 DIV (left) and 15 to 56 DIV (right) for two individual networks cultured in four-nodal mMEAs. LEC LII; lateral entorhinal cortex layer II. DG; dentate gyrus. CA3/CA1; Cornu Ammonis 3/1.

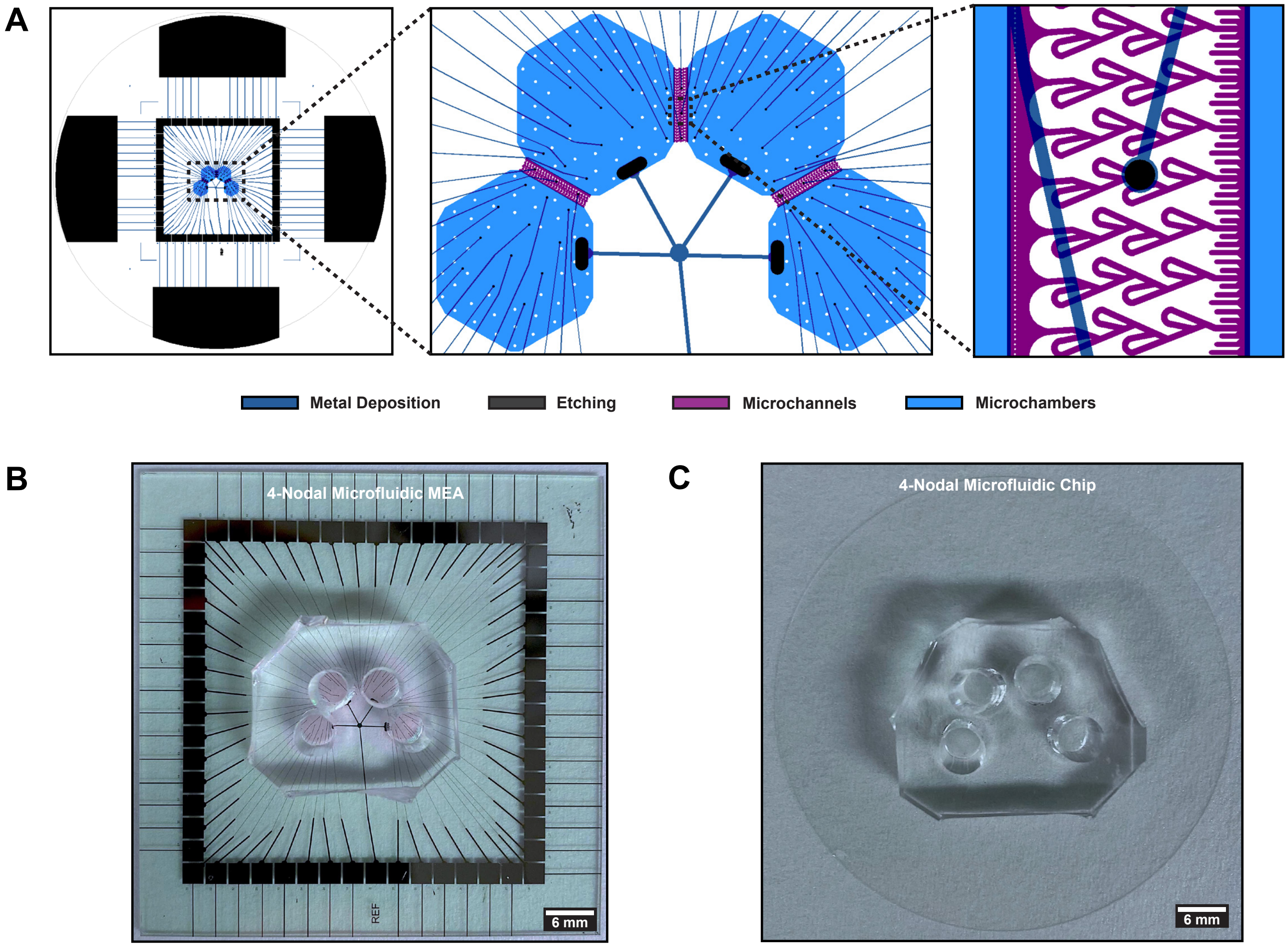


**Figure S5** | **CAD design of the Microfluidic MEAs. (A.)** The microfluidic design with $350\,\mu\text{m}$ long Tesla valve microtunnels. Layer 1 represents the design for the metallization (i.e. forming the microelectrodes and corresponding contact pads), layer 2 the etch mask for etching through the passivation layer, layer 3 the mold for the microfluidic tunnels and layer 4 the mold for the cell compartments. **(B.)** A four-nodal microfluidic chip interfaced with a microelectrode array for electrophysiology. **(C.)** A four-nodal microfluidic chip interfaced with a glass slide, used for immunocytochemistry and fluorescence imaging.

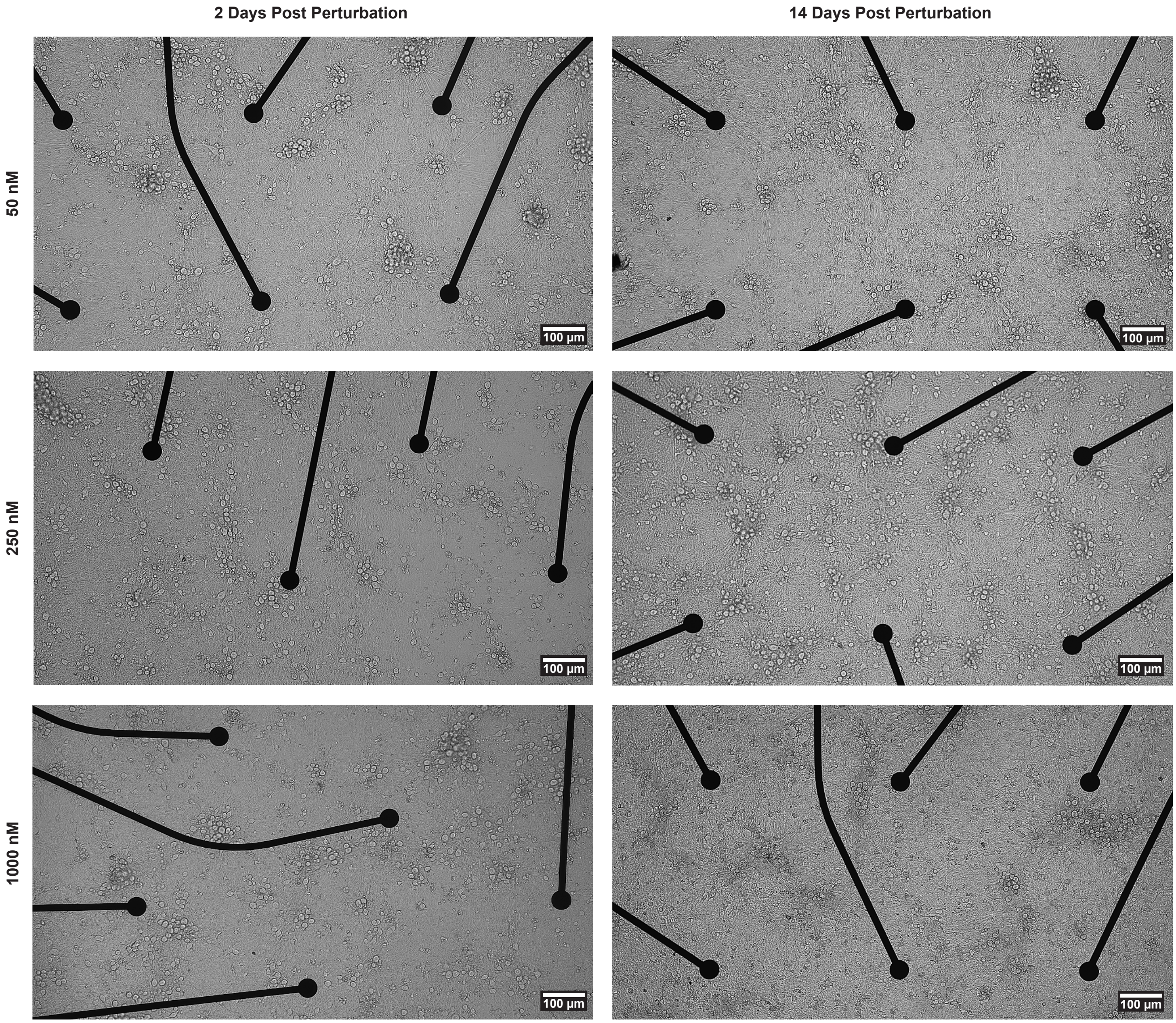


**Figure S6 | Impact of amyloid beta perturbations on network viability.** Concentrations of $50\,\mathrm{nM}$, $250\,\mathrm{nM}$ and $1000\,\mathrm{nM}$ A$\beta$-fragments were applied to neural networks to assess the impact on neural network viability over a period of two weeks. Neither of the concentrations had significant effects on cell viability.

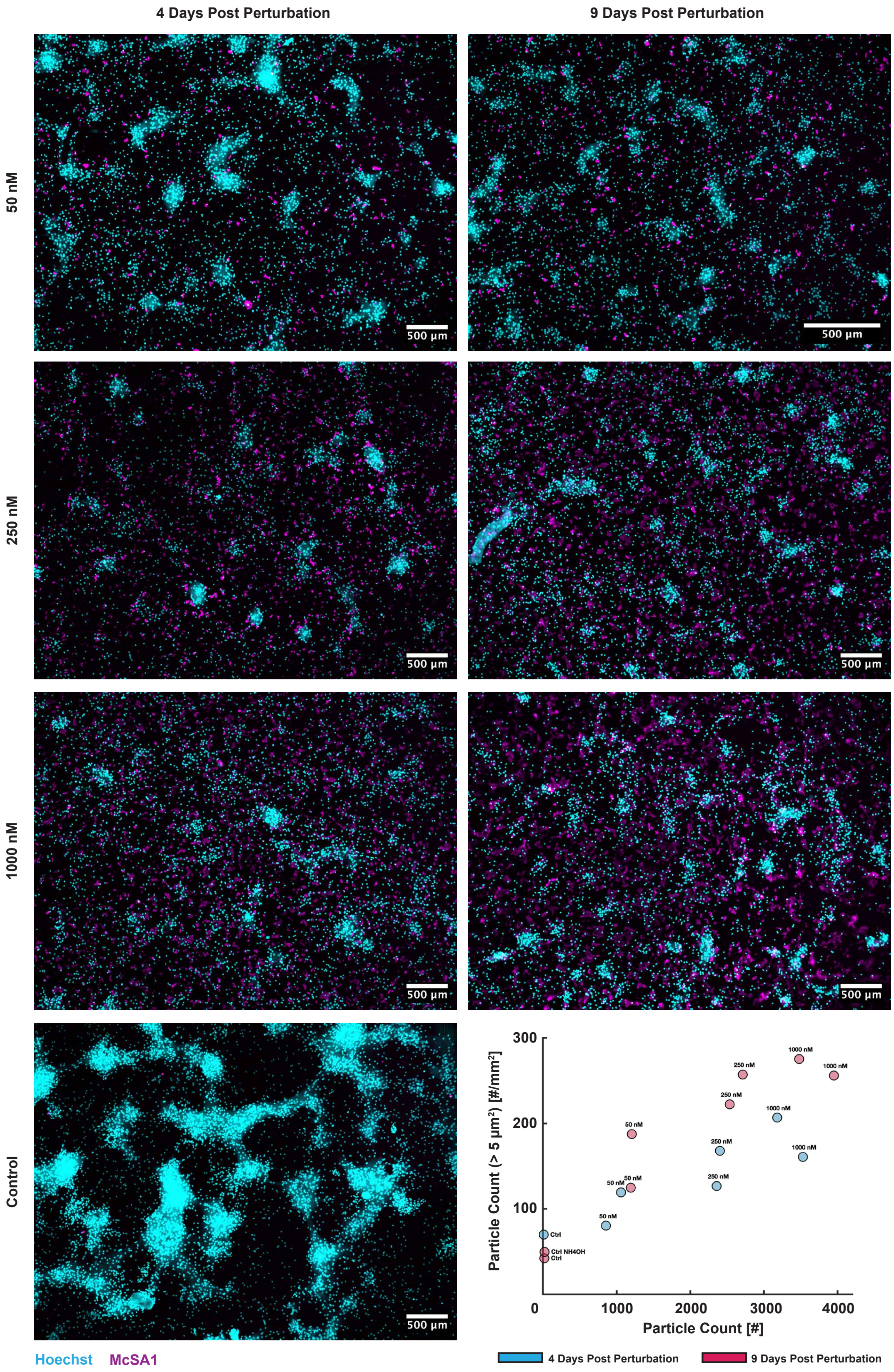


**Figure S7 | Impact of applied amyloid beta concentrations on oligomer formation.** Concentrations of $50\,\mathrm{nM}$, $250\,\mathrm{nM}$ and $1000\,\mathrm{nM}$ A$\beta$-fragments were applied to the neural networks to evaluate their impact on A$\beta$ oligomerisation. Cells were stained for the antibody McSA1, labelling insoluble A$\beta$-aggregates, 4 and 9 days after the fragments were applied. As can be seen from the graph in the lower right corner, increased concentrations of applied A$\beta$-fragments increased both the particle count and average particle size of oligomers in the networks with time.
